# Recombination facilitates adaptive evolution in rhizobial soil bacteria

**DOI:** 10.1101/2021.01.20.427438

**Authors:** Maria Izabel A. Cavassim, Stig U. Andersen, Thomas Bataillon, Mikkel Heide Schierup

**Author notes:** Department of Ecology and Evolutionary Biology, University of California, Los Angeles, 90095, United States (Maria Izabel A. Cavassim). **Corresponding authors**: Maria Izabel A. Cavassim, Mikkel H. Schierup. **Author Contributions Conceptualization**: M.I.A.C., T.B., and M.H.S.; **Methodology**: M.I.A.C., T.B., and M.H.S.; **Formal Analysis**: M.I.A.C.; **Investigation**: M.I.A.C., T.B., and M.H.S.; **Resources**: S.U.A., and M.H.S.; **Data curation**: M.I.A.C., T.B., and M.H.S.; **Writing - Original Draft**: M.I.A.C.; **Writing - Review and Editing**: M.I.A.C., S.U.A., T.B., and M.H.S.; **Visualization**: M.I.A.C., T.B, and M.H.S. **Supervision**: T.B. and M.H.S.; **Project administration**: S.U.A. and M.H.S.; **Funding acquisition**: S.U.A. and M.H.S. **Competing interest** The authors declare that they have no competing interests.

## Abstract

Homologous recombination is expected to increase natural selection efficacy by decoupling the fate of beneficial and deleterious mutations and by readily creating new combinations of beneficial alleles. Here, we investigate how the proportion of amino acid substitutions fixed by adaptive evolution (*α*) depends on the recombination rate in bacteria. We analyze 3086 core protein-coding sequences from 196 genomes belonging to five closely-related species of the genus *Rhizobium*. These genes are found in all species and do not display any signs of introgression between species. We estimate *α* using the site frequency spectrum (SFS) and divergence data for all pairs of species. We evaluate the impact of recombination within each species by dividing genes into three equally sized recombination classes based on their average level of intragenic linkage disequilibrium. We find that *α* varies from 0.07 to 0.39 across species and is positively correlated with the level of recombination. This is both due to a higher estimated rate of adaptive evolution and a lower estimated rate of non-adaptive evolution, suggesting that recombination both increases the fixation probability of advantageous variants and decreases the probability of fixation of deleterious variants. Our results demonstrate that homologous recombination facilitates adaptive evolution measured by *α* in the core genome of prokaryote species in agreement with studies in eukaryotes.

**Significance statement:** Whether intraspecific homologous recombination has a net beneficial or detrimental effect on adaptive evolution is largely unexplored in natural bacterial populations. We address this question by evaluating polymorphism and divergence data across the core genomes of 196 bacterial sequences––belonging to five closely related species of the genus *Rhizobium*. We show that the proportion of amino acid changes fixed due to adaptive evolution (*α*) increases with an increased recombination rate. This correlation is observed both in the interspecies and intraspecific comparisons. By using a population genetics approach our results demonstrate that homologous recombination directly impacts the efficacy of natural selection in the core genome of prokaryotes, as previously reported in eukaryotes.

## Introduction

Genetic recombination is expected to facilitate adaptive evolution by increasing the fixation probability of adaptive mutations and decreasing the probability of fixation of deleterious mutations (McVean and Charlesworth 2000). This is because recombination decouples the fate of adaptive and deleterious variants, decreasing the amount of selective interference throughout the genome (Hill and Robertson 1966; Felsenstein 1974). Selective interference—also termed the Hill-Robertson (HR) effect—is, therefore, strongest in regions of the genome where recombination is low (McVean and Charlesworth 2000). The HR effect is predicted to cause: (i) a reduction in the number of neutral polymorphisms, (ii) the accumulation of slightly deleterious polymorphisms, and (iii) a decrease in the probability of fixation of advantageous alleles (see (Charlesworth et al. 2009)). By mitigating the HR effect, homologous recombination is expected to increase the percentage of amino acid substitutions that are due to adaptive evolution (*α*).

The parameter *α* can also be viewed as the relative proportion between the rate of amino acid changes fixed by positive selection (⍵_*a*_) and the rate of non-adaptive amino acid changes relative to neutral (⍵_*na*_) (as: *α* = ⍵_*a*_/ (⍵_*a*_ + ⍵_*na*_); (Galtier 2016; Moutinho et al. 2020)). Distinguishing between ⍵_*a*_ and ⍵_*na*_ allows us to test more precisely two expectations of the effect of increased homologous recombination: overall genes with higher recombination rates should experience more efficient purifying selection (and hence lower ⍵_*na*_) and increased probability of fixation for beneficial mutations (a higher ⍵_*a*_).

Empirical evidence based on population genomics data supports theory with a positive correlation between recombination and *α* reported in diverse species of eukaryotes, including flies (*Drosophila melanogaster*, (Campos et al. 2014; Castellano et al. 2016)), fungi (*Zymoseptoria tritici* (Grandaubert et al. 2019)), plants (*Arabidopsis thaliana* (Moutinho et al. 2019)), and non-model animal species (Galtier 2016; Moutinho et al. 2020).

Whereas recombination is ubiquitous and mandatory for the reproductive success of most eukaryotes (Page and Hawley 2003), this is not the case for prokaryotes. Nevertheless, many studied prokaryotes show high rates of genetic exchange (Didelot and Maiden 2010) and it is therefore of interest to explore whether such recombination also facilitates adaptive evolution in prokaryotes. Here we study how rates of adaptive evolution and intraspecific homologous recombination co-vary in a species complex of *Rhizobium leguminosarum* responsible for nitrogen fixation in white clover (*Trifolium repens*) nodules. We have previously reported the full genomic sequence of 196 isolates (Cavassim et al. 2020). Of 22115 orthologous gene groups identified among the 196 strains, 4204 genes are present in all isolates (the core genome). Although substantial adaptive evolution might be attributed to accessory genes through their gains and losses via horizontal gene transfer (HGT) (Young et al. 2006; Tian et al. 2010; Porter et al. 2017; Cavassim et al. 2020), we focus here on genes vertically inherited and examine how much variation in rates of homologous recombination explains variation in *α* and its components (⍵_*a*_ and ⍵_*na*_).

Our previous analyses (Cavassim et al. 2020) showed that the 196 strains cluster into five closely related species (2-5% average nucleotide divergence), with horizontal gene transfer between these species only affecting the nitrogen fixation genes and a few well-defined genomic regions––that we exclude in the present analysis. This species complex thus offers a unique opportunity among prokaryotes to estimate the rates of fixation of amino acid changes by adaptive evolution from isolates sampled from natural populations––enabling multiple comparisons of polymorphism and divergence patterns among species. Our analyses provide evidence that the rate of adaptive protein evolution increases with the recombination rate in this species complex.

## Results & Discussion

To estimate the proportion of adaptive evolution (*α*) across this *Rhizobium* species complex and study how *α* covaries with intraspecific recombination rate estimates, we restricted analyses to polymorphism data from regions of the core genome without evidence of recent interspecies HGT.

Of the eighteen species observed within this species complex (Young et al. 2021), we have collected genomic data for five species (gsA-gE). Across all five species (196 strains, gsA:32, gsB:32, gsC:112, gsD:5, gsE:11) (**Fig. S1**), a total of 22115 orthologous gene groups were previously identified (Cavassim et al. 2020); of those, 4204 genes are present in all strains (core genes). Most core genes are found in the large chromosome (3304 genes), but some are located in the chromids (Rh01, Rh02) and one of the plasmids (Rh03) (see Harrison, et al., 2010, and Cavassim et al. 2020). The chromosome, chromids, and the plasmid are hereafter referred to as genomic compartments. To assess the effect of intraspecies homologous recombination on adaptive evolution using a high-quality dataset we filtered out genes that showed evidence of recent interspecies HGT or unexpectedly high rates of nucleotide diversity (see Methods) (**Fig. S2**), leaving a total of 3086 genes (total alignment length: 3091179) and 334040 variable sites for analysis (**Fig. S3**).

First, we estimated nucleotide diversity, intragenic linkage disequilibrium (LD), and the site frequency spectrum (SFS) (see Methods) within each species (**Fig. 1a-c**). The average nucleotide diversity, *π*, an estimator of *2N*_*e*_*μ* in haploids, is significantly different among genomic compartments (**Fig. 1a**, **Table S1**). Across the species, *π* differs by up to a factor of 4.5 (gsA: 0.018, gsB:0.0045, gsC:0.0140, gsD:0.00512, gsE:0.008), with the most polymorphic species being gsA and the least gsB. If we assume similar mutation rates among these closely related species, nucleotide diversity differences reflect interspecies differences in long-term effective population size, *N*_*e*_.

**Figure 1.**
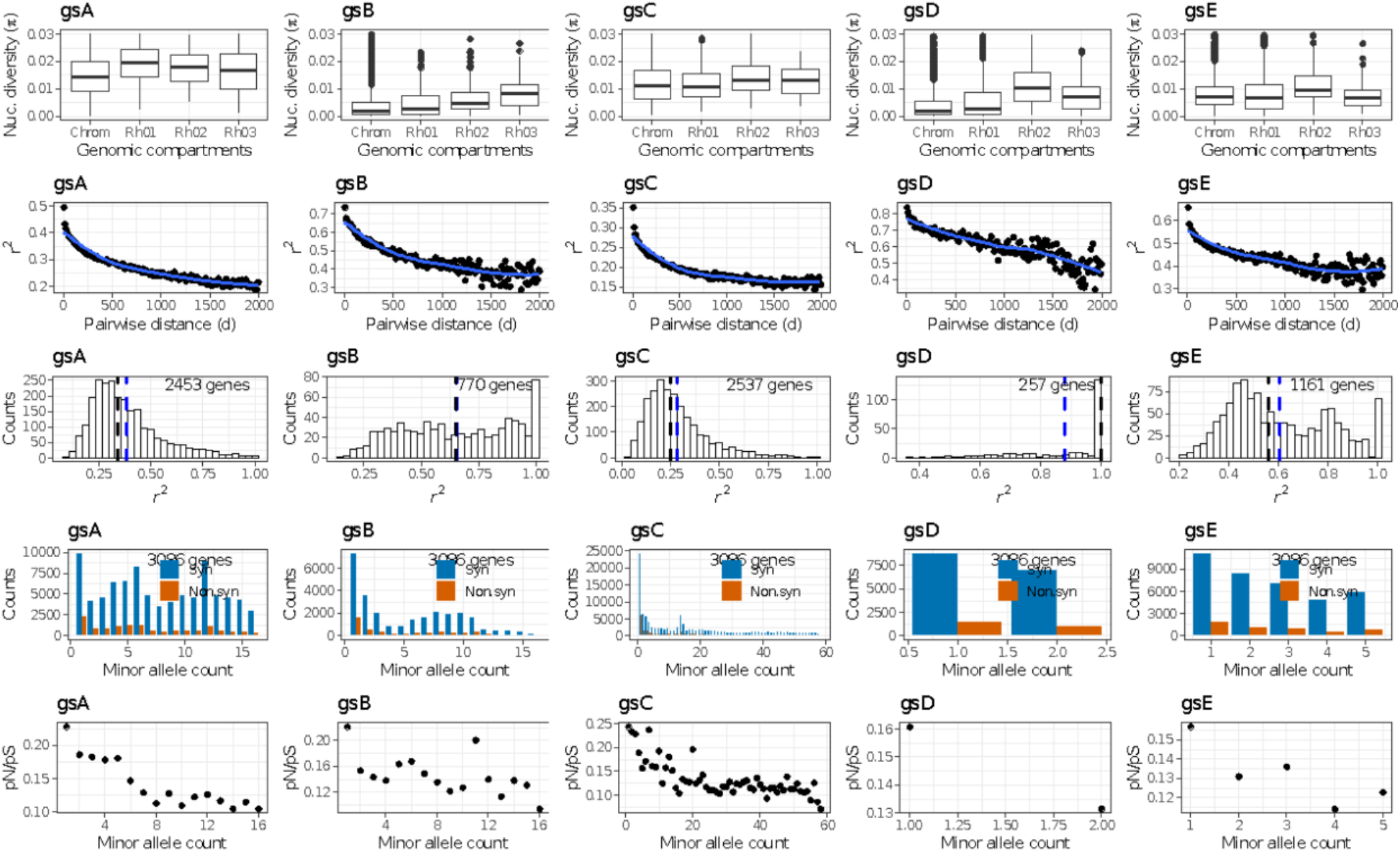
Population genetics parameters across five species. (**a**) Nucleotide diversity (*π*) across 3086 genes distributed along with genomic compartments (chromosome, chromids: Rh01, Rh02, and plasmid: Rh03). To exclude outliers only genes with *π* ≤ *0*.*03* are shown. (**b**) Intragenic linkage disequilibrium measured via the decay of *r*^*2*^ for all core genes (3086 genes). The curve fitting line (in blue) is from a local regression method (loess). (**c**) Linkage disequilibrium (*r*^*2*^) distribution across genes. Only genes with at least ten segregating sites were kept and singletons were excluded (gsA:2453 genes, gsB:770, gsC:2537, gsD:257, and gsE:1161). The black and blue dashed lines correspond to the median and mean *r*^*2*^, respectively. (**d**) Site frequency spectrum counts of synonymous and non-synonymous sites by minor allele count based on all core genes (3086 genes). (**e**) The ratio of non-synonymous to synonymous polymorphisms by minor allele count.

When recombination occurs, we expect that levels of non-random association between pairs of alleles, quantified by measures such as *r*^*2*^ (see Methods), decay with genomic distance (LD decay). To evaluate the recombination rate differences among the five species, we used within-species polymorphism data and computed the average intragenic LD decay for each gene in each species. We observed a rapid decay of LD within the first 1000 base pairs for all species, suggesting substantial amounts of within-species homologous recombination (**Fig. 1b**). The slower decay observed in species gsB either reflects a lower per generation recombination rate or a smaller effective population size (*N*_*e*_). The latter is consistent with the low level of nucleotide diversity measured in gsB. To reliably estimate interspecies differences in *r*^*2*^, we used genes with at least ten informative sites within each species while also excluding variants only found in one strain (singletons) and evaluated their *r*^*2*^ distributions separately (**Fig. 1c**). As expected, the species with the most striking LD decay (gsC) has the lowest *r*^*2*^ (median *r*^*2*^: 0.248), and the opposite is also true (gsD, median *r*^*2*^: 1.00). In summary, these species can be ranked by their recombination levels, from the most recombining to the least, as follows: gsC (median *r*^*2*^: 0.248) > gsA (median *r*^*2*^: 0.341) > gsE (median *r*^*2*^: 0.561) > gsB (median *r*^*2*^: 0.651) > gsD (median *r*^*2*^: 1.00).

Next, we computed the folded site frequency spectrum (SFS) of synonymous and nonsynonymous mutations within each species. Overall, both synonymous and nonsynonymous SFSs differ from the “L” shaped patterns (many rare alleles and fewer frequent alleles) expected in a stationary population at mutation-selection-drift equilibrium (**Fig. 1d**). The observed excess of intermediate frequency SNPs indicates the presence of population structure in some of the species. The effect of population structure is particularly evident in gsC, and this excess is likely driven by strains isolated from French soils (**Fig. S4**). Differences among species suggest distinct demographic histories, with gsC showing an SFS compatible with population expansion and gsA with population decline (Pool et al. 2010).

Using the counts of polymorphism in synonymous and nonsynonymous SFS within each species, we can estimate the overall strength of purifying selection via *pi*_*N*_/*pi*_*S*_. The strength of purifying selection ranks species similarly to their average recombination rate, with more recombining species showing stronger purifying selection (individual *pi*_*N*_/*pi*_*S*_ sorted by recombination rate are: gsC = 0.037, gsA = 0.039, gsE = 0.051, gsB = 0.057, gsD = 0.07). This observation is in line with the theoretical expectation of a positive effect of recombination on the overall efficacy of natural selection. We also observed an excess of rare nonsynonymous relative to synonymous variants (**Fig. 1e**), consistent with the segregation of nonsynonymous variants under weak purifying selection (Ohta 1976). Rare nonsynonymous variants are often deleterious (s~1/*N_e_*) (Hughes et al. 2003; Hughes 2005). Because deleterious variants contribute substantially to polymorphism but rarely to divergence (Fay et al. 2001; Charlesworth and Eyre-Walker 2006), their presence in the genomes, if not controlled for, will lead to an underestimation of *α* (Eyre-Walker and Keightley 2009).

We used GRAPES (Galtier 2016) to estimate the distribution of fitness effects (DFE) (Eyre-Walker and Keightley 2007) and the proportion of adaptive evolution (*α*) from polymorphism and divergence data while accounting for the presence of deleterious mutations. This approach uses the site frequency distribution of both synonymous and nonsynonymous SFS counts to estimate the DFE while accounting for the effect of demography. The significant amount of shared polymorphism among species (**Table S2**) makes it difficult to reliably call ancestral and derived states (Schneider et al. 2011; Tataru et al. 2017). Accordingly, we chose to estimate the DFE and *α* using the folded SFSs (Galtier 2016). To determine the model of the DFE that best fit our data, we used a variety of DFE distribution models (**Table S3**). The DFE models we tested differ by the classes of mutations (deleterious, beneficial, and neutral) included in each DFE model and how fitness effects are distributed within these classes. When using Akaike’s Information Criterion (AIC) to select the best DFE model, we found the GammaZero model overall provides the best fit to the SFS data (**Fig. S5**). This model assumes the existence of weakly deleterious nonsynonymous mutations, modeled as a continuous Gamma distribution (Galtier 2016).

The proportion of adaptive evolution was first computed between all combinations of “mirror” species (*α*_*species1 species2*_), in which “species 2” is used as outgroup (divergence) for “species 1” (polymorphism) and vice-versa (*α*_*species2 species1*_). This yielded twenty combinations in total. Because “mirror” species share an identical history of divergence, their *α* estimates can be considered as “biological replicates” (Galtier 2016) (**Table 1**). Except for the comparison between gsA and gsB, in which differences between *α*_*gsA gsB*_ and *α*_*gsB gsA*_ exceeded 0.09, the overall discrepancy in the values estimated between mirror species does not exceed 0.1. Using each species’ focal polymorphism data, we calculated four *α* estimates by comparing it to the divergence counts of the remaining species (**Table 1**). The most recombining species (gsC) is observed to have the highest *α* across all outgroups used, while the least recombining species (gsD) had the lowest *α* in three out of the four cases.

**Table 1.** The proportion of adaptive evolution (*α*) across pairs of species. The *α* estimates were computed based on the best fitting DFE model (GammaZero) (**Table S3**). For each pairwise estimate of *α* (*α*_*species1 species2*_), the polymorphism data from a focal species (rows) is compared against the divergence counts of an outgroup (columns), and vice-versa (*α*_*species2 species1*_). Confidence intervals are displayed in brackets (grey) and numbers in parentheses represent the *α* ranking (in decreasing order) by outgroup (by column).

| Polymorphism (focal) | Divergence (outgroup) |  |  |  |  |
| --- | --- | --- | --- | --- | --- |
|  | gsA | gsB | gsC | gsD | gsE |
| gsA | - | **0.28**<br>[0.26-0.31] (2) | **0.35**<br>[0.33-0.39] (1) | **0.25**<br>[0.23-0.28] (1) | **0.29**<br>[0.27-0.32] (2) |
| gsB | **0.18**<br>[0.16-0.21] (4) | - | **0.26**<br>[0.24-0.29] (3) | **0.15**<br>[0.13-0.17] (3) | **0.17**<br>[0.15-0.19] (3) |
| gsC | **0.36**<br>[0.33-0.39] (1) | **0.36**<br>[0.33-0.38] (1) | - | **0.25**<br>[0.23-0.28] (1) | **0.30**<br>[0.27-0.32] (1) |
| gsD | **0.25**<br>[0.22-0.28] (3) | **0.25**<br>[0.23-0.27] (4) | **0.25**<br>[0.22-0.28] (4) | - | **0.12**<br>[0.10-0.15] (4) |
| gsE | **0.27**<br>[0.24-0.30] (2) | **0.25**<br>[0.23-0.27] (4) | **0.27**<br>[0.24-0.30] (2) | **0.10**<br>[0.07-0.13] (4) | - |

We then investigated whether intraspecies differences in recombination rate affect the amount of adaptive evolution (*α*) estimated. For each species, we split genes into three recombination classes based on their average *r*^*2*^ values and computed *α* for each class using the GammaZero model (**Fig. S6, Table S4**). Because we only kept genes with at least ten informative sites, the number of genes evaluated across species was different (**Fig. 1c**). For most species comparisons (gsA, gsB, gsC, and gsE), there is a decrease in the proportion of adaptive evolution with a reduction in recombination (increase in *r*^*2*^) (**Fig. 2**). Except for cases in which we used gsD polymorphisms to estimate *α* where all comparisons were non-significant, all the other species pairwise comparisons led to at least one significant difference (based on non-overlapping confidence intervals (CI)) between recombination classes. The low sample size of gsD (5 strains) and its skewed LD distribution (**Fig. 1c**) may have reduced statistical power to discriminate among recombination classes and to estimate *α* reliably.

**Figure 2.**
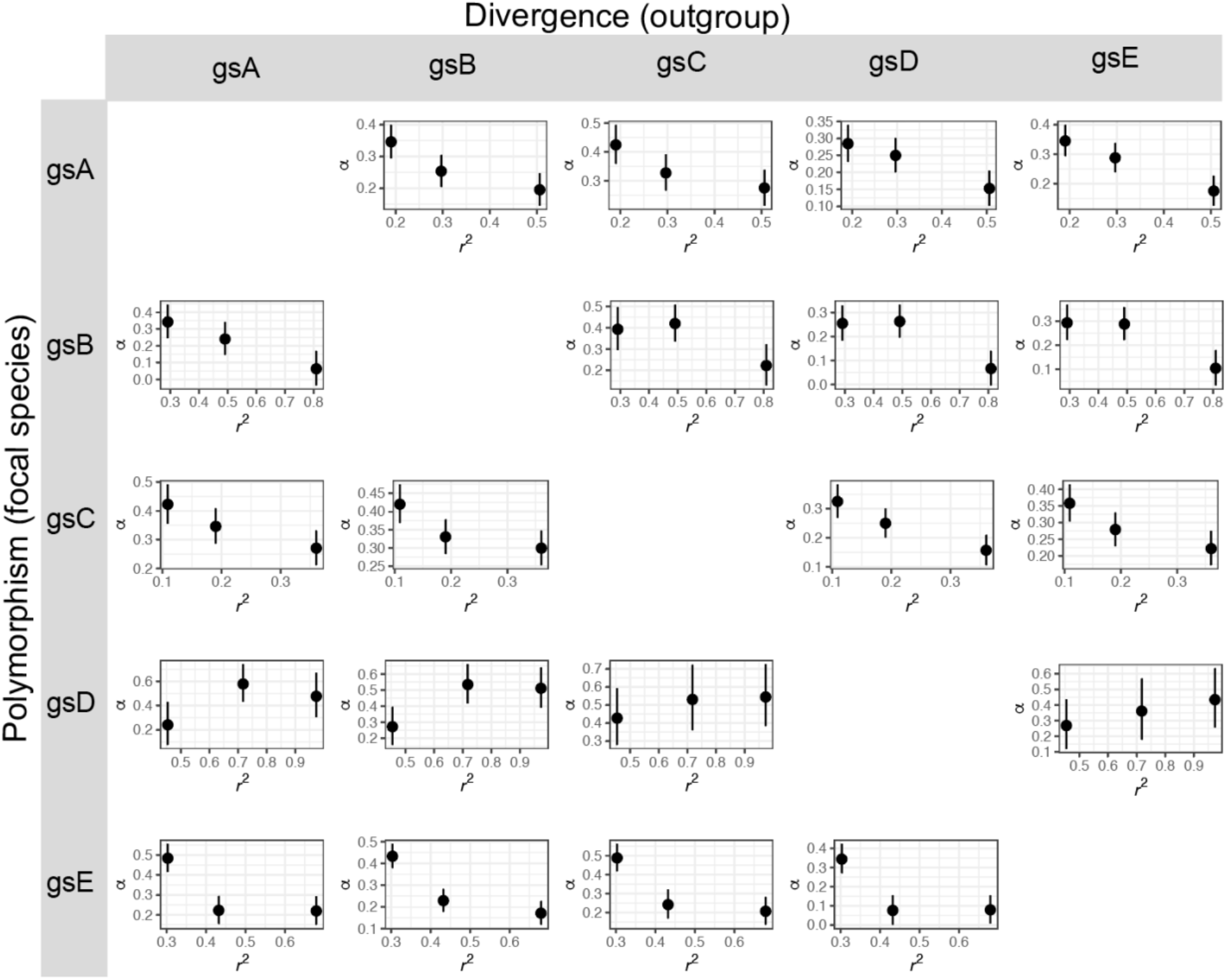
The proportion of adaptive evolution (*α*) by classes of recombination. For each pairwise estimates of *α* the polymorphism data from one species (left in title) is compared against the divergence counts of an outgroup (right in title), and vice-versa. Results are divided into classes of recombination based on *r*^*2*^(a measure that is inversely proportional to the level of recombination). The *α* estimates and their associated confidence intervals were obtained using the best fitting DFE model (GammaZero).

We further assessed the significance of the pattern reported here by permuting, 200 times, across recombination classes (see Methods). Except for simulations in which gsD polymorphisms were used, all the other simulations led to significant differences (p-value ≤ 0.05) among the two most extreme classes of recombination (**Fig. S7**).

The parameter *α* can also be viewed as the relative proportion between the rate of amino acid changes fixed by positive selection (⍵_*a*_) and the rate of non-adaptive amino acid changes (⍵_*na*_): *α* = ⍵_*a*_/ (⍵_*a*_ + ⍵_*na*_). Thus, an increase in *α* with recombination could be due to either an increase in the rate of adaptive substitutions, a decrease in the rate of non-adaptive substitutions, or both. Figure 3 shows that ⍵_*a*_ increases with recombination rate whereas ⍵_*na*_ decreases with recombination rate for most combinations and that the quantitative effects are almost equal in magnitude. Thus, classes of genes evolving under higher recombination rates exhibited lower rates of non-adaptive substitution and increased rates of fixation of adaptive variation. This matches the predictions from selective interference theory (Felsenstein 1974; Barton 1994; McVean and Charlesworth 2000).

**Figure 3.**
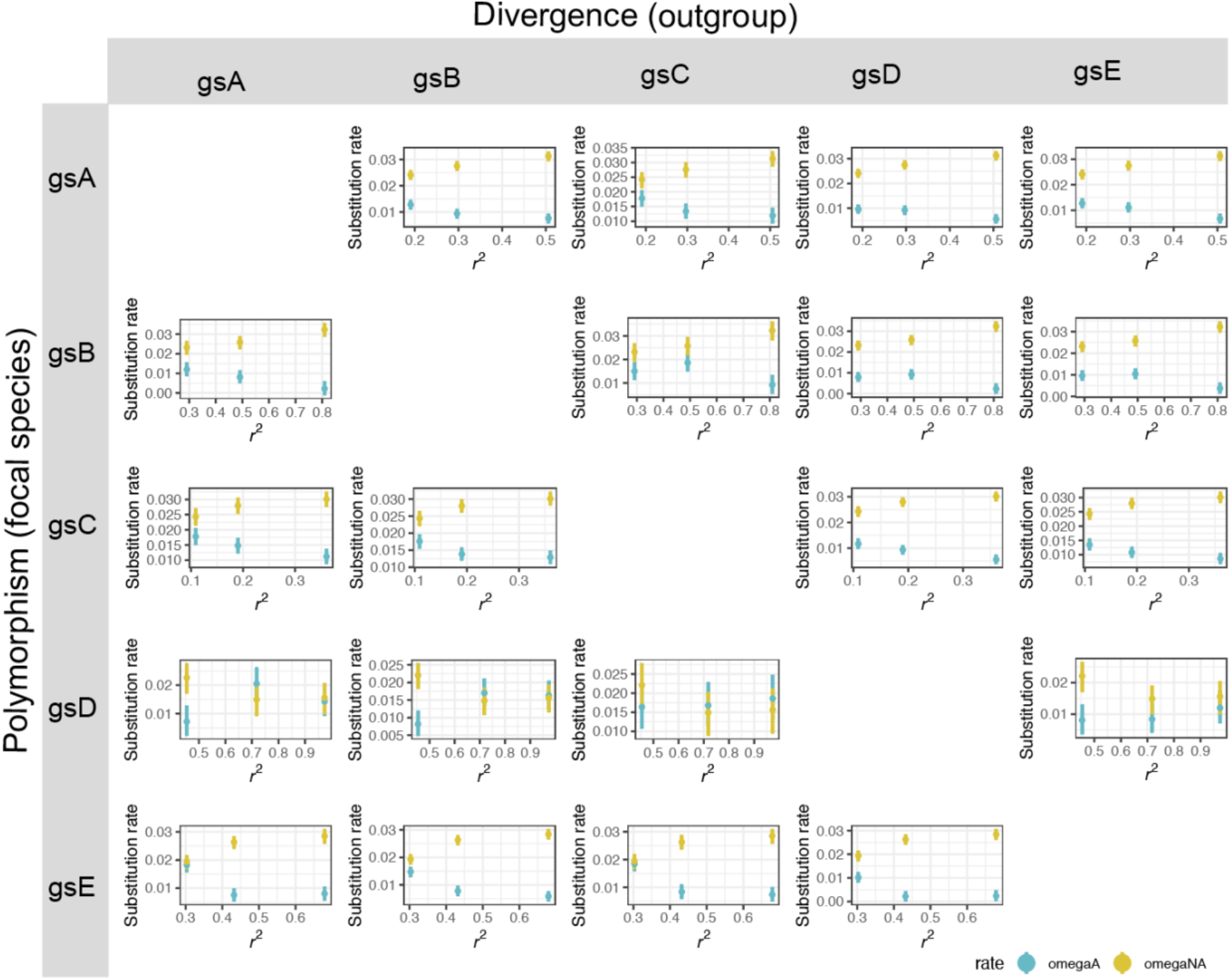
The rates of adaptive (⍵_*A*_) and non-adaptive (⍵_*NA*_) evolution by classes of recombination. For each pairwise estimates of ⍵_*A*_(in blue) and ⍵_*NA*_ (in yellow) the polymorphism data from one species (left in title) is compared against the divergence counts of an outgroup (right in title), and vice-versa. Results are divided into classes of recombination based on *r*^*2*^(a measure that is inversely proportional to the level of recombination). An opposite effect of recombination on ⍵_*A*_ and ⍵_*NA*_ is observed in most pairwise comparisons. The estimates (⍵_*A*_,⍵_*NA*_) and their associated confidence intervals were obtained using the best fitting DFE model (GammaZero).

To evaluate the robustness of these results, we computed two alternative measures of recombination (R/θ, and D’). We then made new recombination classes and evaluated how each recombination measure correlated with *α*. R/θ measures the importance of recombination (R), relative to mutation (θ) across sequences (Vos and Didelot 2009; Didelot and Wilson 2015), while D’ measures LD between sites using a different metric than *r*^*2*^ (see methods). Although the distributions of R/θ and D’ across genes are different than that of *r*^*2*^ (**Fig. 1c, Fig. S8**), these three measures are not independent (Pearson’s correlation between *r*^*2*^and R/θ or D’ ranged from 0.19 to 0.70) (**Fig. S8**). For most species comparisons, the trend between *α* and recombination remains consistent: the higher the amount of recombination (measure by R/θ or D’), the higher *α* is (**Fig. S9–S10**). The mean rates of adaptive evolution (⍵_*a*_) obtained among classes based on D’ or on R/θ estimates also generally agree with those estimated using *r*^*2*^ (**Fig. S11–12**).

The genomes of the Rhizobium species comprise three genomic compartments (chromosomes, chromids, and plasmids) that may have undergone different selection regimes. We tested that by building distinct SFSs for each genomic compartment (see Methods). The mean *a* estimate differs slightly among genomic compartments with higher *a* observed in core genes sampled from the chromosome and the smallest plasmid (Rh03) (**Table 2**), however, the sampling variance of estimates is large, and differences observed are not statistically significant. We also evaluated the recombination rate among these genomic units (**Fig. S13**), recombination is heterogeneous within and across species, as observed for nucleotide diversity (**Fig 1a**). We conclude that we are underpowered to detect differences in the effects of adaptive evolution as a function of recombination between these genomic compartments.

**Table 2.** The proportion of adaptive evolution (*α*) across genomic compartments. The *α* estimates were computed based on the best fitting DFE model (GammaZero) (**Table S3**). For each genomic compartment (chrom = chromosome; Rh01 and Rh02 = chromids; Rh03 = plasmid) we compared the polymorphism data from species gsA against the divergence counts of an outgroup (rows). Confidence intervals are displayed in brackets (grey).

| Species | Genomic compartments |  |  |  |
| --- | --- | --- | --- | --- |
|  | Chrom | Rh01 | Rh02 | Rh03 |
| gsA | - | - | - | - |
| gsB | 0.4393<br>[0.29-0.60] | 0.3593<br>[0.20-0.53] | 0.2192<br>[0.02-0.44] | 0.4484<br>[0.26-0.66] |
| gsC | 0.4399<br>[0.24-0.67] | 0.3116<br>[0.10-0.56] | 0.2413<br>[-0.03-0.57] | 0.4682 [0.26-0.71] |
| gsD | 0.3568<br>[0.19-0.54] | 0.3247<br>[0.17-0.50] | 0.1383<br>[-0.09-0.41] | 0.3444<br>[0.16-0.56] |
| gsE | 0.4013<br>[0.24-0.58] | 0.3650<br>[0.22-0.52] | 0.2235<br>[0.03-0.45] | 0.4203<br>[0.25-0.61] |

In this study, we applied a methodology that estimates the proportion of amino acid changes that have been fixed by positive selection (*α*) while also estimating the individual components of *α* (⍵_*a*_ and ⍵_*na*_). We have only included genes that are present in all sampled genomes––the so-called core genome. Because the methodology used requires summarizing the data by building an SFS, the analyses cannot be readily extended to the accessory genome–– given that many accessory genes are a result of HGT (Popa and Dagan 2011), a DFE computed from this source will not necessarily reflect the DFE of the studied species. The accessory genome represents roughly 40% of the genome of these species (Cavassim et al., 2020), and likely also contributes to adaptive evolution (e.g., acquired symbiotic ability (Cavassim et al., 2020; Kumar et al., 2016)) as also shown in other studies (e.g., antibiotic resistance in *Staphylococcus aureus* (Harris et al. 2010); or adaptation to a new ecological niche (Ochman et al. 2000; Wiedenbeck and Cohan 2011)).

It has been previously shown experimentally that both recombination via plasmid-mediated gene transfer (conjugation) or via transformation can accelerate bacterial adaptation in populations of *E. coli* (Cooper 2007) and *Helicobacter pylori* (Baltrus et al. 2008). These studies are in line with previous simulation studies (Cohen et al. 2005; Levin and Cornejo 2009). Homologous recombination was shown to accelerate adaptation when incorporated in simulations, including mutation and selection, at rates typical of species like *E. coli*, *H. influenzae*, *B. subtilis* (Levin and Cornejo 2009). Cohen et al., 2005 studied recombination in the context of a simple fitness landscape model with evolution implemented as a continuous Markov process and observed a drastic speed up on the rate of adaptive evolution with increased population sizes. Cooper 2007 concluded that recombination only had a positive effect on adaptation when beneficial mutations were abundant in the population––implying that standing genetic variation, possibly driven by higher mutation rate (*μ*) or higher effective population size (*N*_*e*_), may be crucial for recombination to be useful. Assuming that mutation rate (*μ*) is similar among these sibling species, then the observed adaptive differences among them may reflect differences in effective population size, recombination rate, or a combination of both (Cohen et al. 2005; Arnold et al. 2018).

## Conclusion

We have found that five bacterial species within the species complex *R. leguminosarum* display different yet high levels of recombination. The estimates of *α* ranged between 0.07 and 0.39 among species. These estimates are lower than those based on 410 orthologs observed in *E. coli* (0.58, CI=0.45, 0.68) but close to estimates from *S. enterica* (0.34, CI=0.14, 0.50) previously reported (Charlesworth and Eyre-Walker 2006)––however, in this study population fluctuations were not accounted for.

Levels of recombination correlate—both across and within species—with higher amounts of adaptive evolution estimated either as the rate of adaptive substitutions (⍵_*a*_) or as the proportion of amino acid changes that have been fixed by positive selection (*α*). For instance, the most recombining species (gsC) consistently exhibited the largest *α*, independent of the outgroup used. Within each species, we also find a positive correlation between intragenomic recombination rate and *α*. This is both due to a higher estimated rate of adaptive evolution (⍵_*a*_) and a lower estimated rate of non-adaptive evolution (⍵_*na*_), suggesting that recombination both increases the fixation probability of advantageous variants and decreases the probability of fixation of deleterious variants. These findings are robust to the measure of recombination (*r*^*2*^, *R*/*θ*, and D’) used to define classes and the choice of outgroup used for computing divergence. Despite variation in recombination rate among genomic compartments, we did not observe significant differences in adaptive evolution among them.

The positive association between amounts of homologous recombination on *α* we report here is in line with population genetic studies conducted in vertebrates (Galtier 2016; Moutinho et al. 2020) and invertebrates (Presgraves 2005; Betancourt et al. 2009; Arguello et al. 2010; Mackay et al. 2012; Campos et al. 2014; Grandaubert et al. 2019); it is also in line with experimental and simulation studies of adaptive evolution in prokaryotes (Cohen et al. 2005; Cooper 2007; Baltrus et al. 2008; Levin and Cornejo 2009). It points to recombination being a general facilitator of adaptive evolution across the tree of life.

## Material and methods

### Identification of orthologous genes

We previously isolated and sequenced 196 strains from white clover (*Trifolium repens*) root nodules harvested in Denmark, France, and the UK. To identify a set of orthologous genes shared across strains, we followed the methods outlined in Cavassim et al., 2020. Briefly, the strains were previously subjected to whole-genome shotgun sequencing using 2×250bp Illumina paired-end reads (Illumina, USA). Genomes were assembled using SPAdes (v. 3.6.2, (Bankevich et al. 2012)) and assembled further, one strain at a time, using a custom Python script (Jigome, available at https://github.com/izabelcavassim/Rhizobium_analysis/tree/master/Jigome).

From the assembled genomes (Cavassim et al. 2020), we predicted protein-coding sequences using prokka (Seemann 2014) (v1.12); this resulted in a total of 1468264 protein-coding sequences. To predict orthologous genes from these sequences, we used Proteinortho (Lechner et al. 2014; Seemann 2014) (v5.16b) with default parameters except for enabling the synteny flag. We identified a total of 22115 orthologous gene groups, including a total of 17911 orthologous observed in at least two strains (accessory genes), and 4204 orthologous found in all 196 strains (core genes).

Orthologous gene groups were aligned using clustalo (Sievers et al. 2011) (v.1.2.0) in a codon-aware manner. To determine the genetic relationship among all 196 strains, we previously calculated their pairwise average nucleotide identity (ANI) across 305 conserved orthologous gene alignments (Cavassim et al. 2020). Under the 95% ANI threshold that delineates species boundaries (Konstantinidis et al. 2006), we demonstrated that these 196 Rhizobium strains constitute five distinct *R. leguminosarum* species (gsA, gsB, gsC, gsD, and gsE) (**Fig. S1**). To ensure that we had a high-quality orthologous dataset for extracting segregating sites, we filtered it further (see below).

### Filtering out orthologous gene groups with evidence of interspecies HGT or misassigned orthologous gene groups

To evaluate the impact of intraspecific homologous recombination on adaptive evolution, we excluded genes that showed signals of recent HGT across the five species analyzed. We have previously developed and applied a phylogenetic method to quantify HGT (introgression score) (Cavassim et al. 2020). This method evaluates the possible number of shifts from one species to another in a given phylogenetic tree. The pipeline takes a gene tree as an input and traverses the tree––using the depth-first search approach––searching deeper in the tree whenever possible. Once the tip is reached the species classification for that given strain is stored. A list containing the species in order of search is collected for the entire tree, the introgression score is then computed as the number of shifts from one species to another in the list minus the set of species plus 1.

We previously showed that most of the core genes shared among the present species respect the species-tree topology (introgression score = 0) (Cavassim et al. 2020). The exceptions are genes sitting in the symbiosis conjugative plasmids, and two chromosomal islands (introgression score > 7). To ensure that we were only analyzing high-quality gene alignments, with no evidence of misassigned orthologous gene groups and with little evidence of HGT, we imposed some restrictions. We only accepted genes that passed the following criteria: (i) were present in every strain (196 strains), (ii) with a nucleotide diversity (*π*) below 0.1 (see **Fig. S2a**), (iii) identifiable replicon origin (chromosome and chromids), (iiii) and with an introgression score ≤ 3. A total of 3086 out of 4204 core genes were kept, and of these, 2550 genes were found in the chromosome, 288 genes in chromid Rh01, 160 genes in chromid Rh02, and 88 genes in plasmid Rh03.

### Variant calling

To identify single nucleotide polymorphisms (SNPs) along with our high-quality set of core genes, we evaluated each gene codon-aware alignment using a custom python script https://github.com/izabelcavassim/Popgen_bacteria. For a given core gene alignment and position, we first counted the number of unique nucleotides (A, C, T, G). Only sites containing two unique nucleotides were considered variable sites (bi-allelic SNPs). SNP matrices were then built and encoded as follows: major alleles were encoded as 1 and minor alleles as 0. The nucleotide diversity (*π*), gene length, and the distributions of segregating sites across core genes are described in **Fig. S2(b-d)**.

### Transition transversion rate bias (kappa) and expected number of synonymous and nonsynonymous sites

Because transitions are more often synonymous at third codon positions than are transversions, to correctly identify the expected number of synonymous (*Lps*) and nonsynonymous sites (*Lpn*), we first estimated the average transition/transversion rate bias (kappa) (Ina 1995) across species. To this end, we followed the methods described in (Yang and Nielsen 2000) and used two classes of sites: fourfold-degenerate sites at the third codon positions and nondegenerate sites. Mutations at the fourfold-degenerate sites are synonymous, and therefore kappa at those sites should reflect only the mutational bias. All mutations at non-degenerate sites are nonsynonymous and were also used to estimate kappa. We computed an average kappa by combining these two classes based on equations 8, 9, 10, and 11 of (Yang and Nielsen 2000). These equations have been implemented within the CodonSeq class in Biopython (Cock et al. 2009) (private function “_count_site_YN00()”), and these private functions were adapted to our dataset.

To estimate a common kappa for each gene alignment (including all species and strains), we averaged estimates from pairwise analyses across 50 randomly chosen strains. The kappa distribution has a mean of 5.6 and a median of 5.20 (**Fig. S3a**), we used the median to compute the expected number of synonymous and nonsynonymous sites. To this end, we followed the methods described by (Ina 1995) and modified by (Yang and Nielsen 2000)––also implemented within Biopython. A total of 284742 synonymous, 49298 nonsynonymous sites were counted (**Fig. S3b-c**).

### Divergence sites and shared polymorphisms

For each pair of species (a focal and an outgroup), we evaluated their variable sites and computed the number of shared synonymous (*pS*) and nonsynonymous (*pN*) polymorphisms. Given a bi-allelic SNP (0 and 1), we considered shared polymorphic sites as sites for which both alleles (0,1) were segregating in both species (**Table S2**). We restricted the estimates of divergence to those sites for which we had variable sites across species. We classified synonymous (*d*_*S*_) and nonsynonymous divergent sites (*d*_*N*_) as those sites in which we observed fixed differences between a focal species and an outgroup.

### Calculating the folded Site Frequency Spectrum

One can infer the distribution of fitness effects from Site Frequency Spectrum (SFS) data (Eyre-Walker and Keightley 2009). Because of the amount of shared polymorphism among the present species (**Table S2**), it becomes problematic to confidently distinguish ancestral from derived polymorphisms (Hernandez et al. 2007). Therefore, we chose to estimate the DFE using a method that uses the folded site frequency spectrums (SFS) of synonymous and nonsynonymous sites (Galtier 2016). To this end, we built the folded synonymous and nonsynonymous site frequency spectrums by tabulating the observed counts of the minor allele frequencies. The synonymous and nonsynonymous SFSs, and the divergence counts, were then used to estimate the DFE and the proportion of adaptive substitutions (*α*) across pairs of species.

### Calculating the strength of purifying selection

The strength of purifying selection was measured as the ratio of nucleotide diversity at nonsynonymous (*pi*_*N*_) and synonymous sites (*pi*_*S*_). For each gene and class of polymorphisms (synonymous and nonsynonymous) nucleotide diversity was computed as: 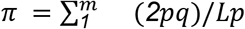, in which *p* and *q* are the allele frequencies, and Lp is the expected number of synonymous (*Lps*) or nonsynonymous positions (*Lpn*) along the gene. We use the median of the *pi*_*N*_/*pi*_*S*_ distribution among genes as a proxy for the strength of purifying selection per species.

### Estimation of adaptive and non-adaptive non-synonymous substitutions rates

Fitted parameters of the DFE were used to compute the expected *d*_*N*_/*d*_*S*_ under the different models, which was compared to the observed *d*_*N*_/*d*_*S*_ to estimate the adaptive substitution rate (*ω*_*A*_); non-adaptive substitution rate (*ω*_*NA*_), and the proportion of adaptive substitutions (*α*) with *ωA* = *α d*_*N*_/*d*_*S*_ and *ω*_*NA*_ = (*1* − *α*) *d*_*N*_/*d*_*S*_.

To account for potential departures of the SFS from demographic equilibrium (assuming the Wright-Fisher model)––possibly driven by changes in the effective population size or by population structure––the method uses nuisance parameters to correct for these SFS distortions (Eyre-Walker et al. 2006). The different DFE models were compared using the Akaike Information Criterion (AIC) (Akaike 1992).

### Recombination rate estimates

To estimate the recombination rate per gene per species, we used three approaches: two based on the degree of association (or linkage disequilibrium) between pairs of alleles in a sample of haplotypes (*r*^*2*^and *D′*), and a third approach, ClonalFrameML (R/θ) (Vos and Didelot 2009; Didelot and Wilson 2015), which relies on the maximum likelihood inference to detect recombination events that disrupt a clonal pattern of inheritance in bacterial genomes.

#### (1) linkage disequilibrium (*r*^2^)

Intragenic linkage disequilibrium (LD) measures the correlation between pairs of alleles with genomic distance in a gene. Here we used Pearson’s *r*^*2*^ correlation measure.

Each gene genotype matrix (containing a minimal set of ten single nucleotide polymorphisms (SNPs)) was first normalized as follow: let *N* denote the total number of individuals, and *M* the total number of SNPs, the full gene genotype matrix (*X*) has dimensions *N* × *M* with genotypes encoded as 0’s and 1’s for the *N* haploid individuals. Each column *S_i_* (*i* = 1, …, *M*) of the *X* matrix is a vector of SNP information of size *N*. To compute LD, we discarded SNPs found only in one sample (singletons). We then applied a Z-score normalization to each SNP vector by subtracting the vector by its mean and dividing it by its standard deviation 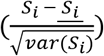, resulting in a vector with mean 0 and variance 1. The linkage disequilibrium was then calculated as a function of distance *d* (maximum 1000 base pairs apart) and was computed as the average LD of pairs of SNPs *d* base pairs away from each other. The calculations were done in the following way:

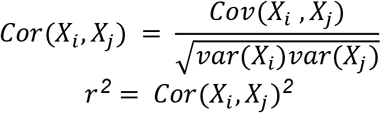

In which *j* > *i* and *X*_*i*_ is composed of the genotypes of all individuals of a given species for SNP position *i* in the genotype matrix. *X*_*j*_is formed of the genotypes of all individuals of the same species for position *j* in the genotype matrix, and*d* = *j* − *i* with *d* ≤ *1000* base pairs. Results were then summarized into bins of 100 base pairs apart; for each bin, a mean *r*^*2*^ was computed and then averaged to a singular *r*^*2*^ value.

#### (2) linkage disequilibrium (D’)

The average linkage disequilibrium within genes was also measured by D’ (Lewontin 1964) as follow: given two locus *i* and *j* (with alleles *A* and *a*, observed in locus *i* and alleles *B* and *b* observed in locus *j*), we first computed the frequency of all possible haplotype combinations (*f*_*AA*_, *f*_*AB*_, *f*_*Ab*_, *f*_*ab*_) and allele frequencies (*f*_*A*_, *f*_*B*_, *f*_*a*_, and *f*_*b*_). The coefficient D was then computed as: *D*_*AB*_ = *f*_*AB*_ − *f*_*A*_ ⋅ *f*_*B*_ and D’ was computed by standardizing |D| by its maximum possible value as: 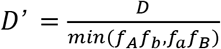, *if D* > *0*, or 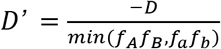, *if D* < *0*. An average value per gene was stored and computed similarly to the *r*^*2*^ statistics (see above).

#### (3) ClonalFrameML

To estimate the changes in the clonal phylogeny by recombination (R), relative to mutation (θ) (R/θ), we used the software ClonalFrameML (Vos and Didelot 2009; Didelot and Wilson 2015). For each species, we first concatenated all core gene alignments (3086 genes) to build the starting phylogenetic species tree using a maximum likelihood approach (Raxml-ng (Kobert et al. 2014; Stamatakis 2014)). We then input each phylogenetic tree within each gene alignment to estimate R/θ.

### Calculating the significance levels between recombination classes

To test whether differences in *α* among recombination classes were statistically significant across species comparisons, we conducted a non-parametric test by shuffling genes among recombination classes (200 permutations) and recording the amplitude of differences between *α* estimates (*Δ*_*α*_ = *max*_*α*_ − *min*_*α*_). We calculated a p-value by comparing the observed *Δ*_*α*_ against the simulated *Δ*_*α*_ distribution.

### Estimation of adaptive substitutions by genomic compartments

To estimate adaptive evolution (*α*) among genomic compartments we first down-sampled the genes from genomic compartments (chromosome, Rh01, Rh02) to reach the size of the smallest genomic compartment (Rh03). Due to the paucity of data, we chose to compute *α* using polymorphism data from the most polymorphic species (gsA) and contrasted it against each outgroup (gsB-gsE). The *α* estimates and their associated confidence intervals were obtained using the GammaZero DFE model within GRAPES.

## Data sharing plans

- Code generated for this study can be found at https://github.com/izabelcavassim/Popgen_bacteria.
- The data that support the findings of this study are available in the INSDC databases under Study/BioProject ID PRJNA510726 (https://www.ncbi.nlm.nih.gov/bioproject/PRJNA510726/).
- Accessions numbers are from SAMN10617942 to SAMN10618137 consecutively.
- Orthologous gene alignments and SNP matrices are available on FigShare (file Data.zip): https://doi.org/10.6084/m9.figshare.11568894.v5

## Acknowledgments

The authors would like to thank industrial partners DLF Trifolium, SEGES, and Legume Technology Ltd. for their contribution to the field trials. The authors would also like to thank J. Peter W. Young, Paula Tataru, Bjarni Vilhjálmsson, and Marjolaine Rousselle for their helpful discussions about this work.

## Supplemetary Information

**Figure S1:**
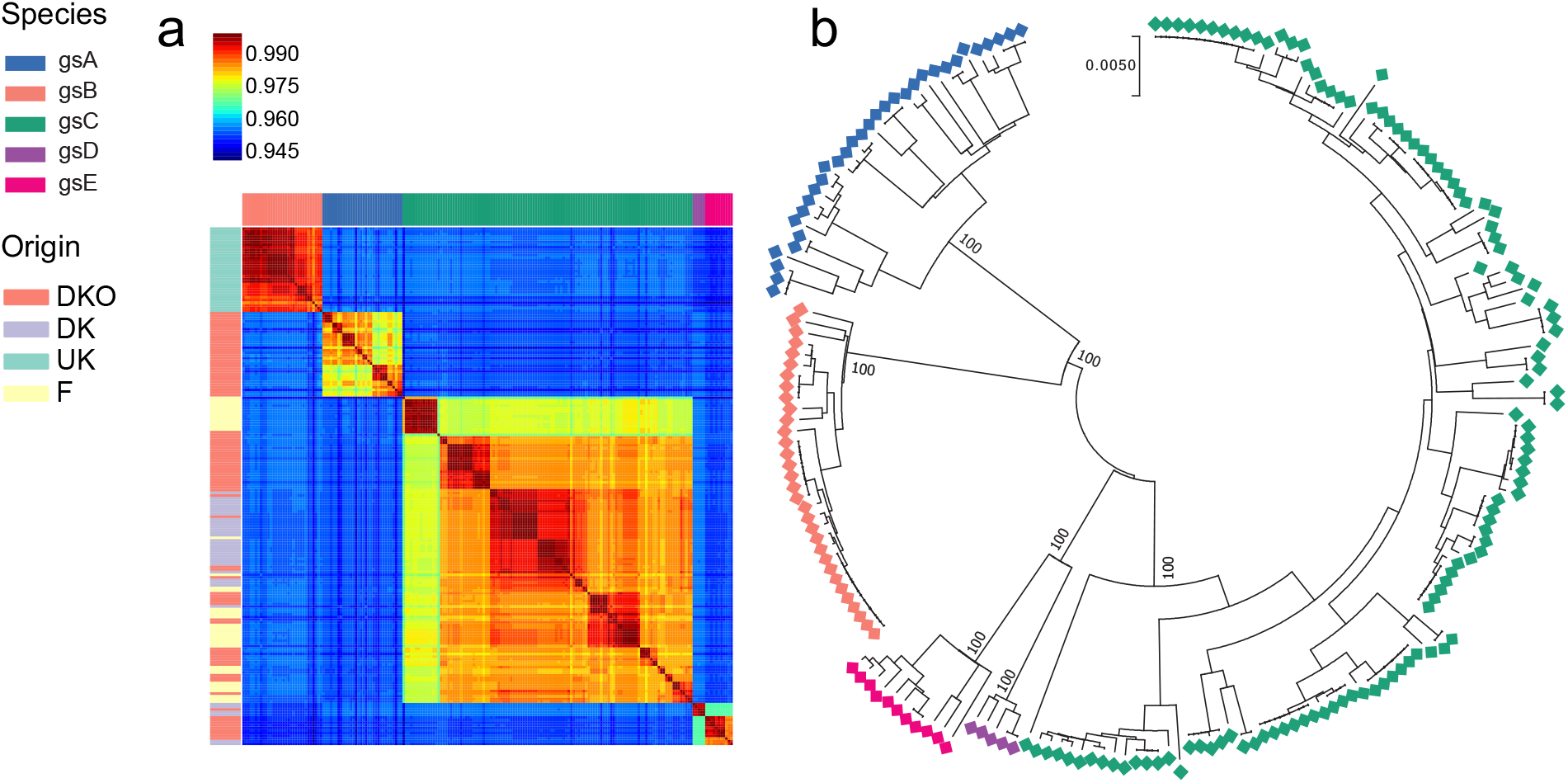
Pairwise average nucleotide identity (ANI) estimates (196×196) for concatenated sequences of 305 housekeeping genes (**Fig. A**, see Cavassim et al., 2020 for details). The top x-axis coloured bars represent the species classification (gsA-gsE) based on 95 percent ANI threshold (Konstantinidis et al. 2006). The y-axis corresponds to the sampling origin of strains and are indicated by coloured bars (Denmark organic fields in red (DKO), Denmark conventional fiels in purple (DK), France in yellow (F), and United Kingdom in green (the UK)). Species phylogeny based on the concatenation of 305 housekeeping genes using the neighbourjoining method (**Fig. B**). Tips are coloured by species (gsA-gsE) and the numbers in nodes indicate the bootstrap value.

**Figure S2:**
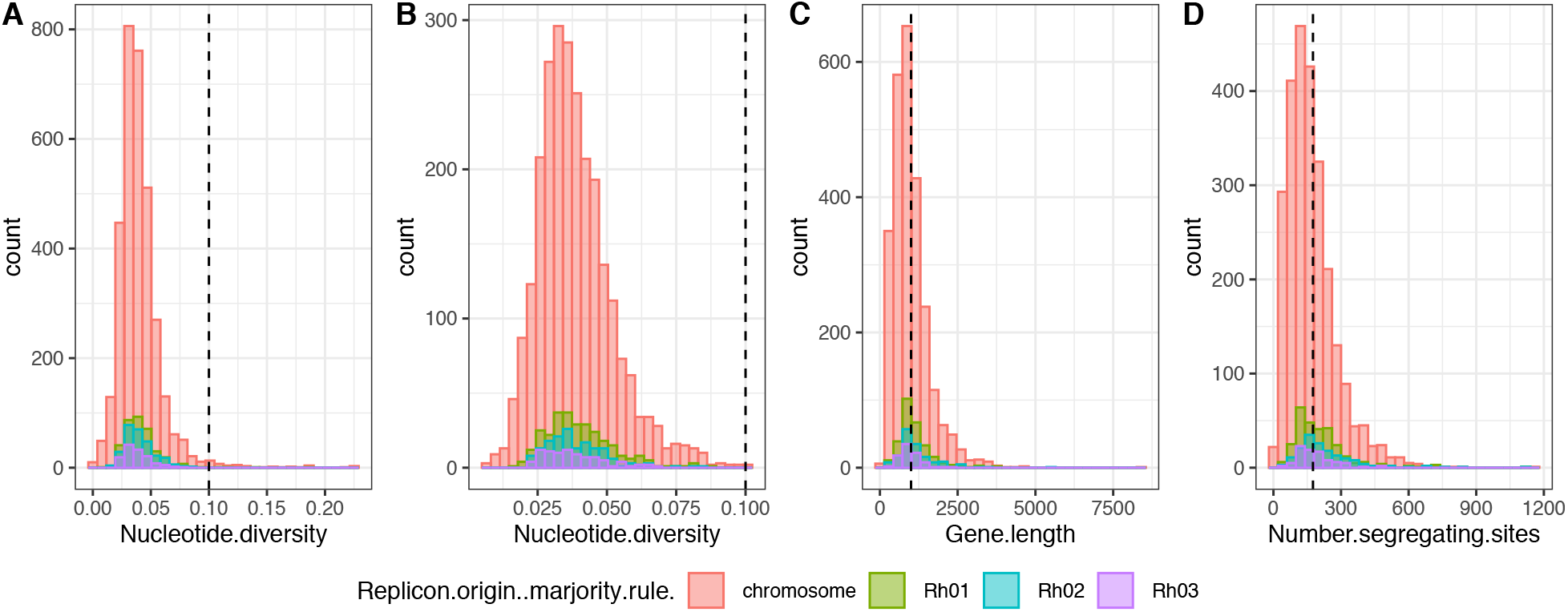
Population genetics statistics of core orthologs. The nucleotide diversity (*π*) distribution, gene length and the number of segregating sites based on a total of 4202 core genes. To compute polymorphism and divergence counts we required high quality gene-alignments with little evidence of horizontal gene transfer. The nucleotide diversity distribution of core genes (**Fig. A**) was used to impose a nucleotide diversity threshold (<=0.10). Other thresholds were also set (see Material and Methods), and a total of 1116 genes were discarded (**Fig. B**). The remaining genes were on average 1000 base pairs long (**Fig. C**) with an average of 175 segregating sites (across all 196 strains) (**Fig. D**).

**Figure S3:**
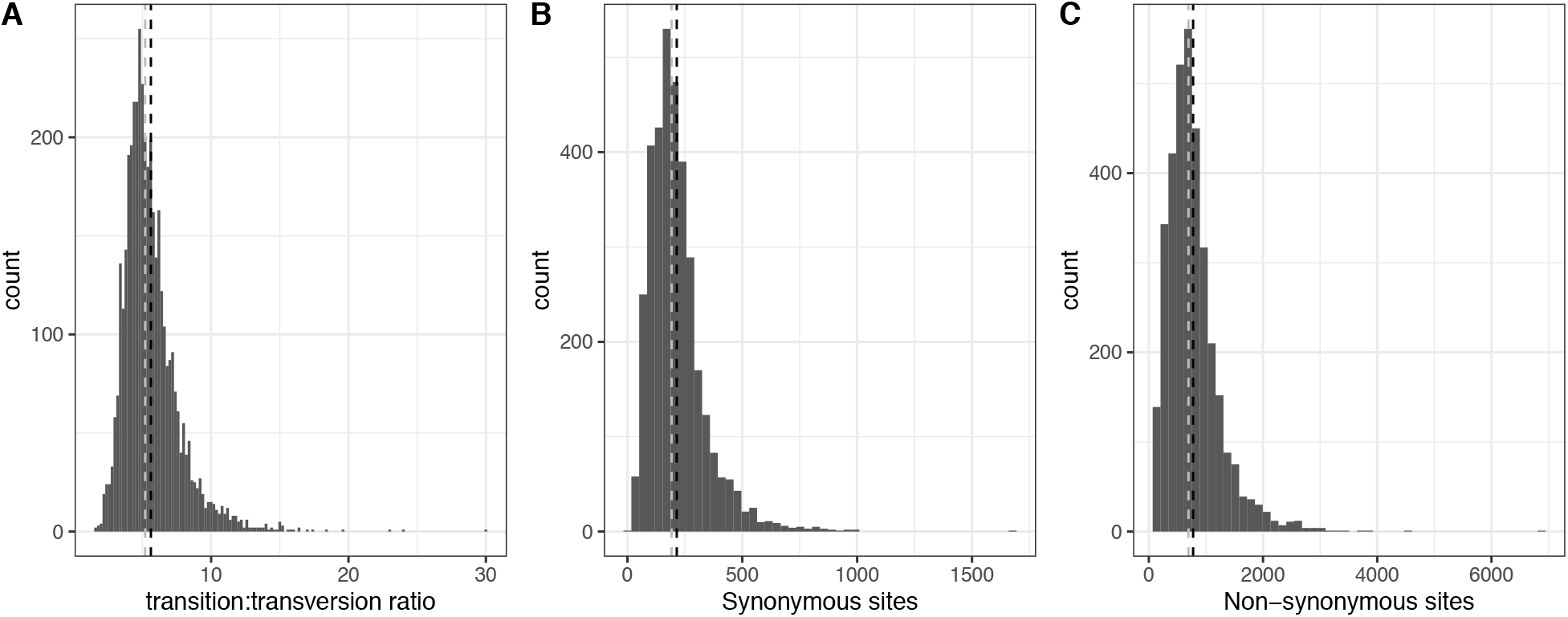
Transition/Transversion (kappa), Synonymous-sites, and Nonsynonymous sites distributions across core genes. To calculate the expected number of synonymous and nonsynonymous sites we first computed the transition/transversion ratio (kappa) across 3086 genes (**Fig. A**). Black and grey dashed lines correspond to the mean and median kappa values, respectively. Synonymous and nonsynonymous sites were computed based on the kappa median (5.20), and the distributions of counts for both site types are shown in **Fig. B-C**.

**Figure S4:**
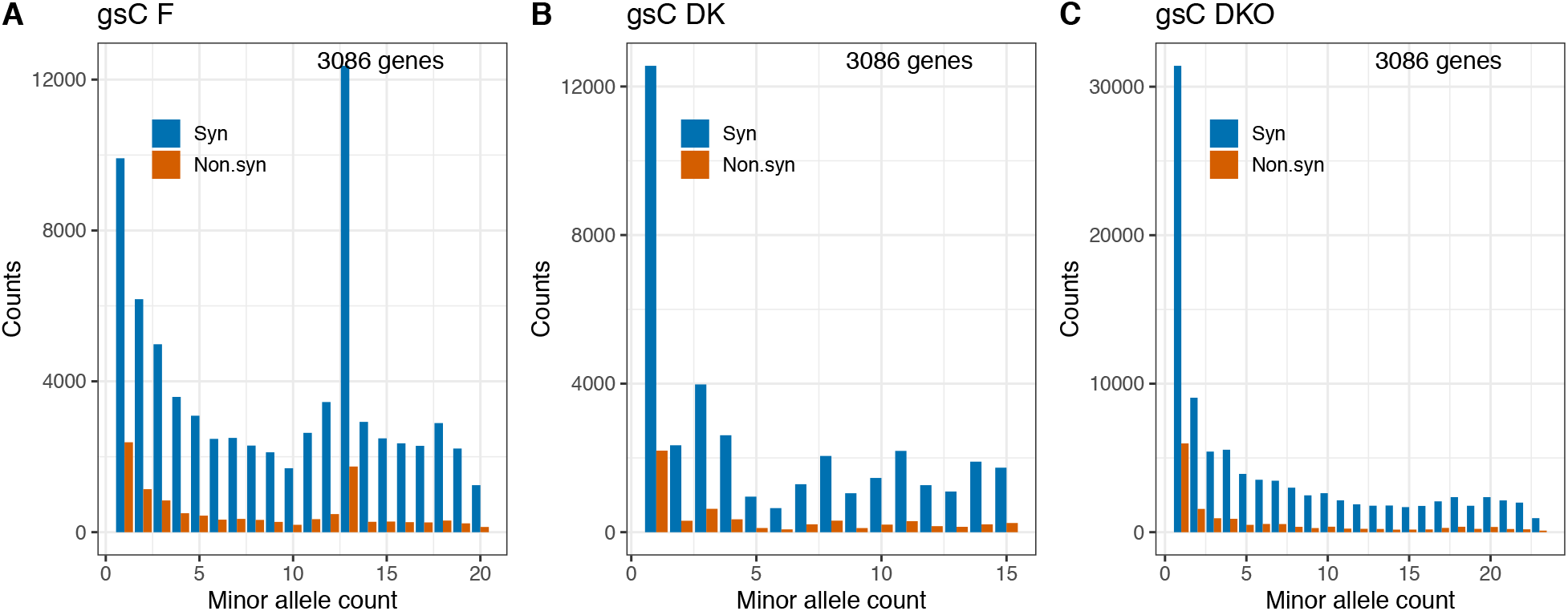
Population structure and the SFS of species C (gsC). Isolates from gsC were found in different countries (France, **Fig. A**, and in Denmark, **Fig. B-C**). The gsC strains found in Denmark were isolated from soils subjected to different agricultural treatments (breeding field trials (gsC DK, **Fig. B**) and organic fields (gsC DKO, **Fig. C**)). The shape of SFS varies across different origins, with an evident deviation from the “L” shaped patterns (many rare alleles and fewer frequent alleles) observed in strains sampled from France (**Fig. A**).

**Table S1:**
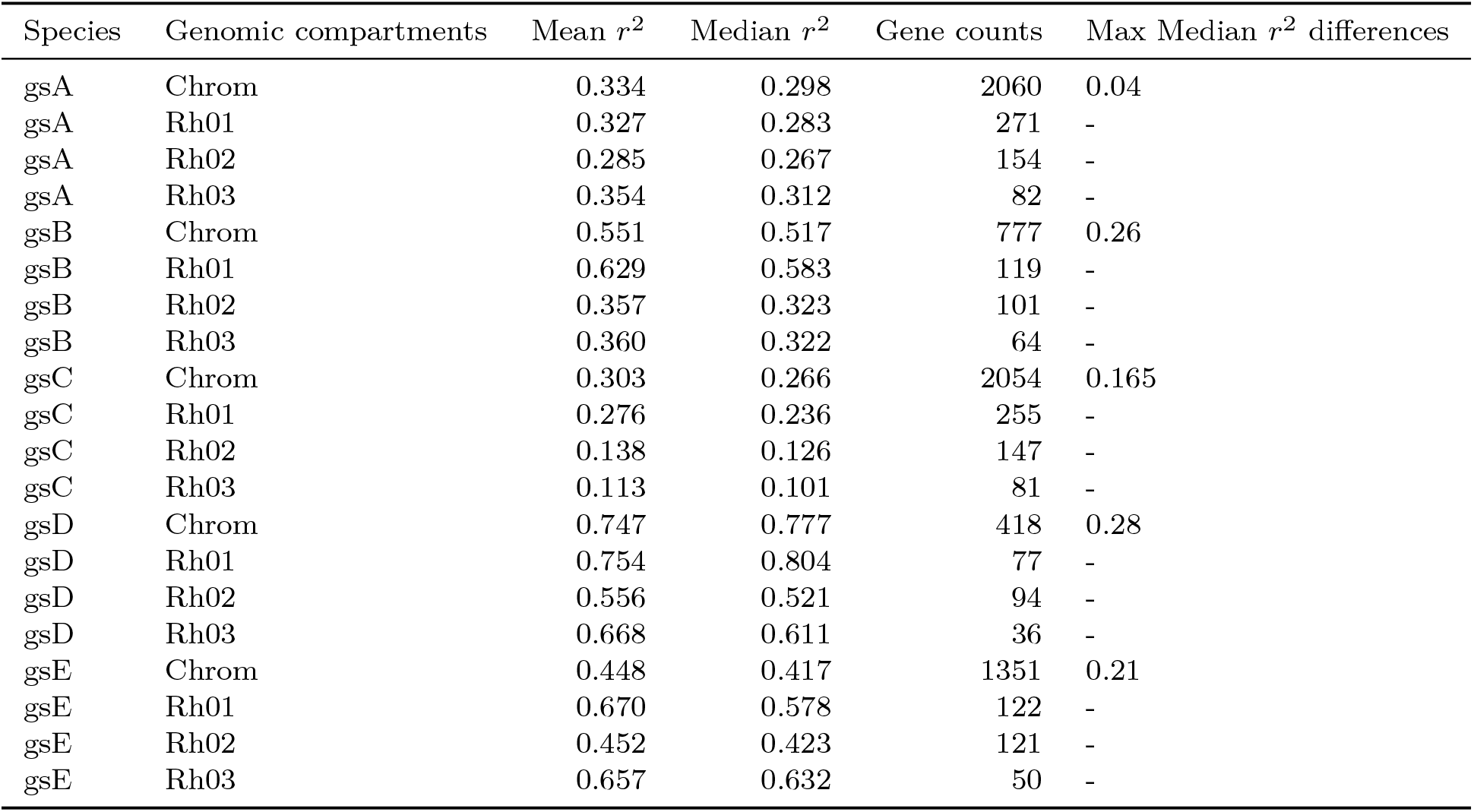
The recombination rate variation across species and their genomic compartments. (chromosome, chromids (Rh01 and Rh02) and plasmid (Rh03)).

| Species | Genomic compartments | Mean $r^2$ | Median $r^2$ | Gene counts | Max Median $r^2$ differences |
| --- | --- | --- | --- | --- | --- |
| gsA | Chrom | 0.334 | 0.298 | 2060 | 0.04 |
| gsA | Rh01 | 0.327 | 0.283 | 271 | - |
| gsA | Rh02 | 0.285 | 0.267 | 154 | - |
| gsA | Rh03 | 0.354 | 0.312 | 82 | - |
| gsB | Chrom | 0.551 | 0.517 | 777 | 0.26 |
| gsB | Rh01 | 0.629 | 0.583 | 119 | - |
| gsB | Rh02 | 0.357 | 0.323 | 101 | - |
| gsB | Rh03 | 0.360 | 0.322 | 64 | - |
| gsC | Chrom | 0.303 | 0.266 | 2054 | 0.165 |
| gsC | Rh01 | 0.276 | 0.236 | 255 | - |
| gsC | Rh02 | 0.138 | 0.126 | 147 | - |
| gsC | Rh03 | 0.113 | 0.101 | 81 | - |
| gsD | Chrom | 0.747 | 0.777 | 418 | 0.28 |
| gsD | Rh01 | 0.754 | 0.804 | 77 | - |
| gsD | Rh02 | 0.556 | 0.521 | 94 | - |
| gsD | Rh03 | 0.668 | 0.611 | 36 | - |
| gsE | Chrom | 0.448 | 0.417 | 1351 | 0.21 |
| gsE | Rh01 | 0.670 | 0.578 | 122 | - |
| gsE | Rh02 | 0.452 | 0.423 | 121 | - |
| gsE | Rh03 | 0.657 | 0.632 | 50 | - |

**Figure S5:**
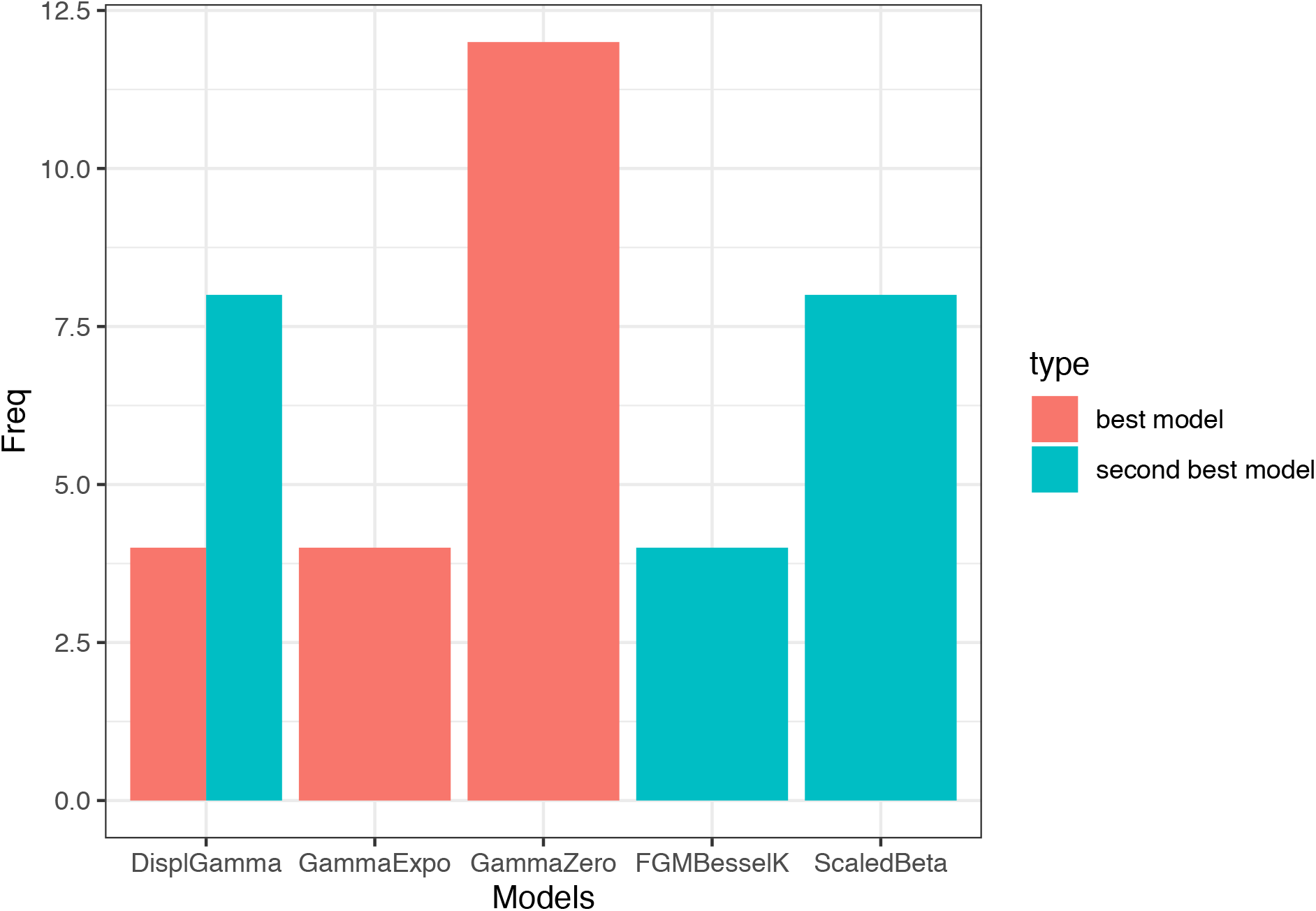
Model selection of DFE models for the between species estimation of *α*. To estimate *α* an assumption of the shape of the distribution of fitness effects (DFE) is required. To determine the model of the DFE that best fit the data, a variety of DFE distribution models were tested, with assumptions described in the Sup. Table 3. For each model and each SFS, the Akaike’s information criterion (AIC) was computed. The best model was defined as the model that had most often the lowest AIC. The frequency of the best and second best models are shown. The GammaZero model was found to be the best model 12 out of 20 times.

**Table S2:** The proportion of shared polymorphisms (‘shared poly’) among species. Pn, and Ps refer to the total number of non-synonymous and synonymous polymorphisms, respectively.

| Species1 | Species2 | Pn (sp1) | Ps (sp1) | Pn (sp2) | Ps (sp2) | Shared non poly | Shared syn poly | Prop shared poly (sp1) | Prop shared poly (sp2) |
| --- | --- | --- | --- | --- | --- | --- | --- | --- | --- |
| gsA | gsB | 12974 | 86783 | 4457 | 26682 | 406 | 5686 | 0.061 | 0.196 |
| gsA | gsC | 12974 | 86783 | 18700 | 107889 | 1774 | 21531 | 0.234 | 0.184 |
| gsA | gsD | 12974 | 86783 | 2325 | 15724 | 239 | 3271 | 0.035 | 0.194 |
| gsA | gsE | 12974 | 86783 | 5069 | 37227 | 412 | 6027 | 0.065 | 0.152 |
| gsB | gsC | 4457 | 26682 | 18700 | 107889 | 543 | 6843 | 0.237 | 0.058 |
| gsB | gsD | 4457 | 26682 | 2325 | 15724 | 61 | 967 | 0.033 | 0.057 |
| gsB | gsE | 4457 | 26682 | 5069 | 37227 | 100 | 1869 | 0.063 | 0.047 |
| gsC | gsD | 18700 | 107889 | 2325 | 15724 | 291 | 3758 | 0.032 | 0.224 |
| gsC | gsE | 18700 | 107889 | 5069 | 37227 | 513 | 7213 | 0.061 | 0.183 |
| gsD | gsE | 2325 | 15724 | 5069 | 37227 | 124 | 1618 | 0.097 | 0.041 |

**Figure S6:**
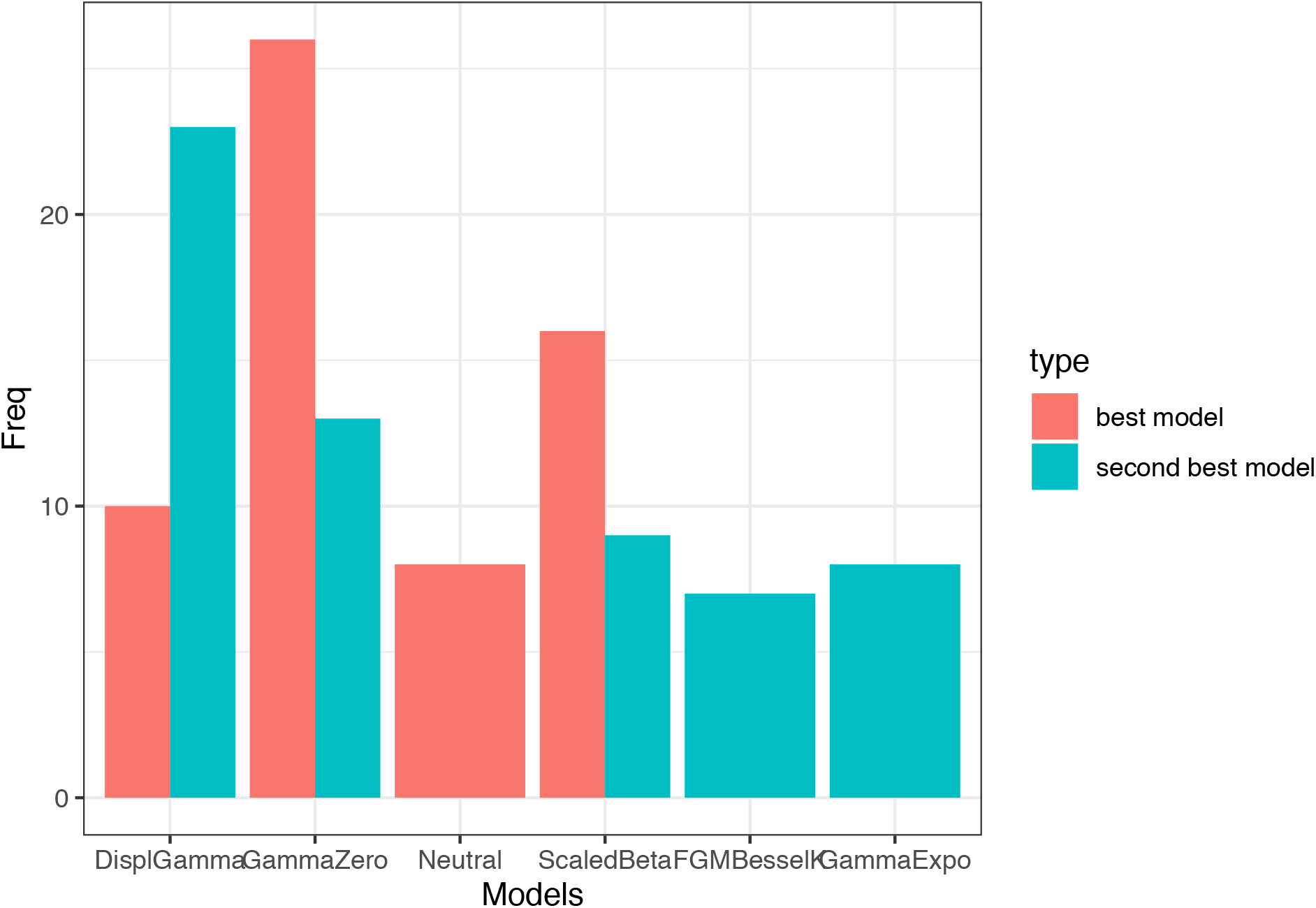
Model selection of DFE for intraespecific estimation of *α*. The Akaike’s information criterion (AIC) of competing models were computed (see **Fig. S4** legend). The GammaZero model was found to be the best model 30 out of 60 times.

**Table S3:**
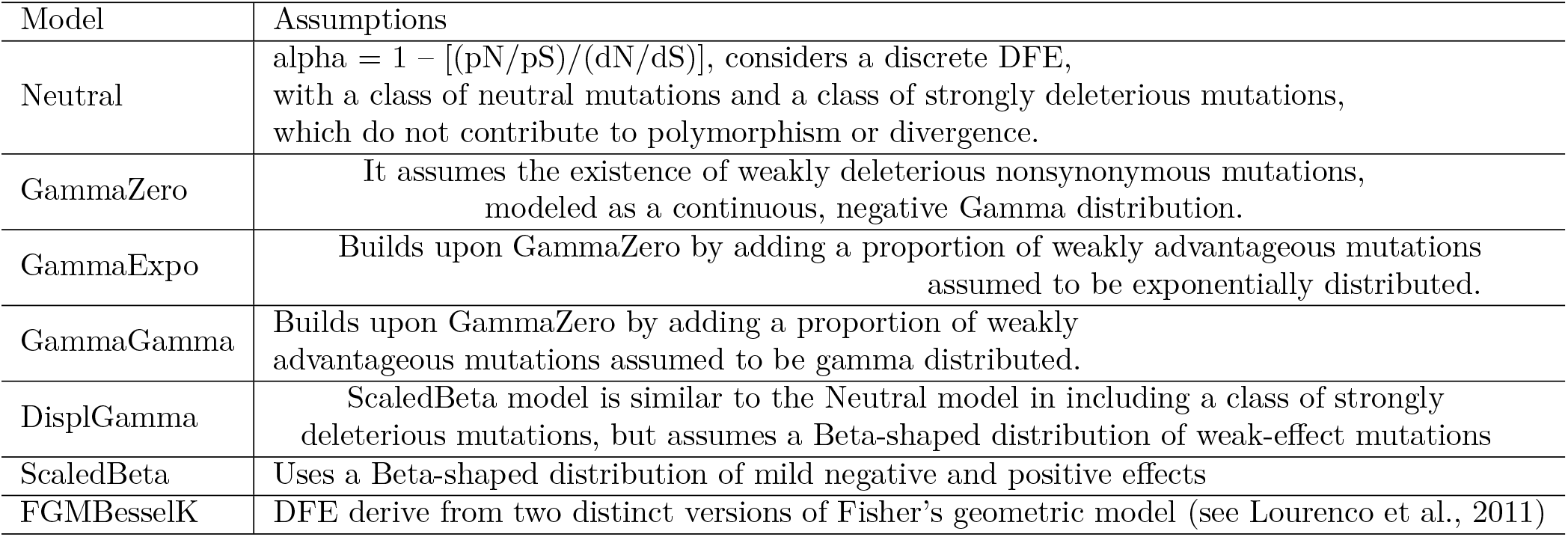
Modelling the distribution of fitness effects (DFE). Description of the models used to compute the distribution of fitness effects and alpha using the software Grapes (Galtier et al., 2016).

**Figure S7:**
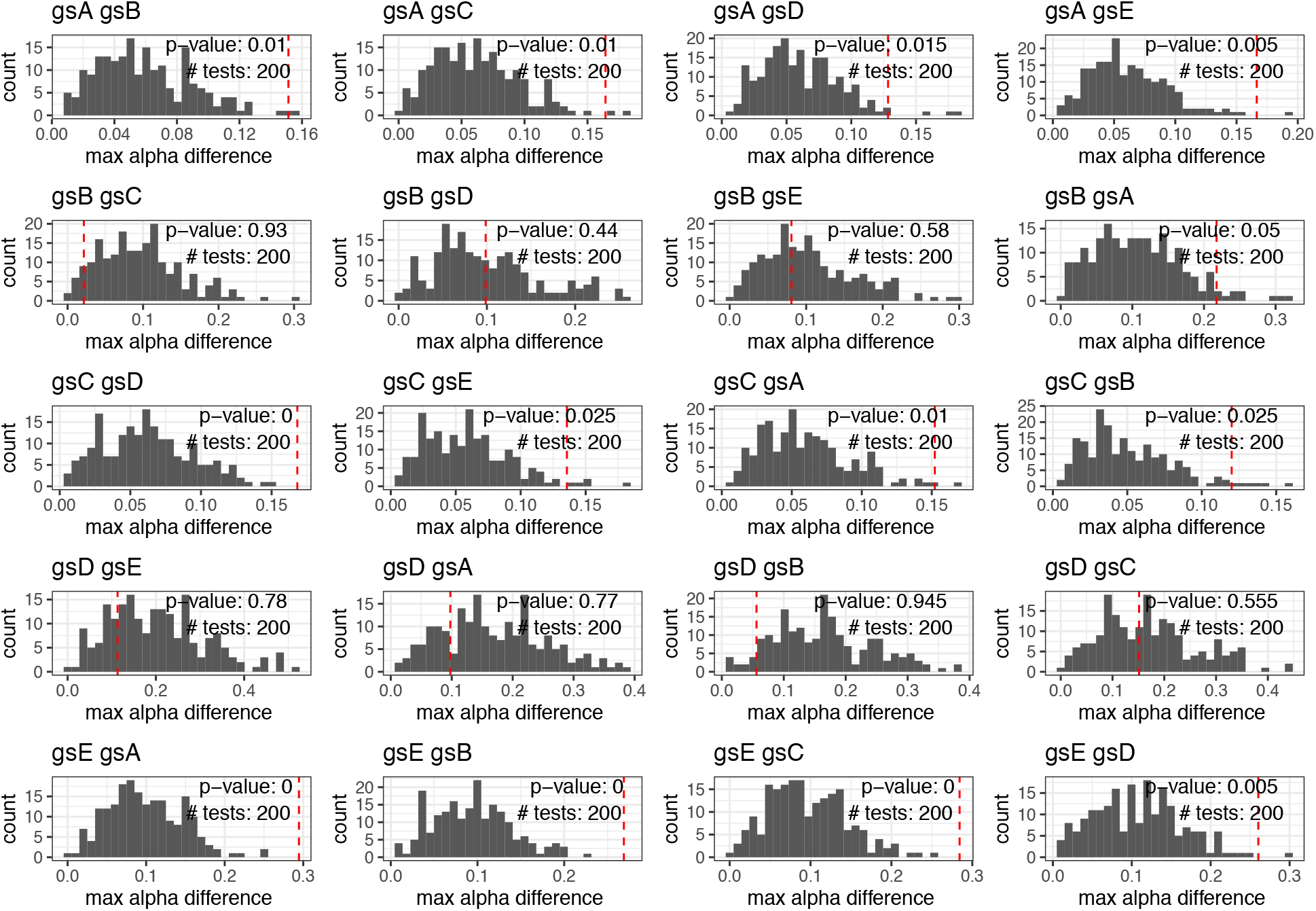
Null distributions of intragenic differences in *α* across recombination classes. For each pairwise estimates of *α* (*α_species_*_1*species*2_), the polymorphism data from a focal species (species 1) is compared against the divergence counts of an outgroup (species 2), and vice-versa (*α_species_*_2*species*1_)). We simulated a null-distribution by permuting the recombination classes of genes (200 times), and for each permuted dataset computing the magnitude of differences between – estimates across recombination classes (Δ_*α*_ = *max_α_* - *min_α_*). The Δ_*α*_s distributions (histograms) were compared against each observed Δ_*α*_ (red dashed line) (see **Fig. 3**). P-values were computed as the number of simulations above or equal to the observed Δ_*α*_ (red dashed lines).

**Figure S8:**
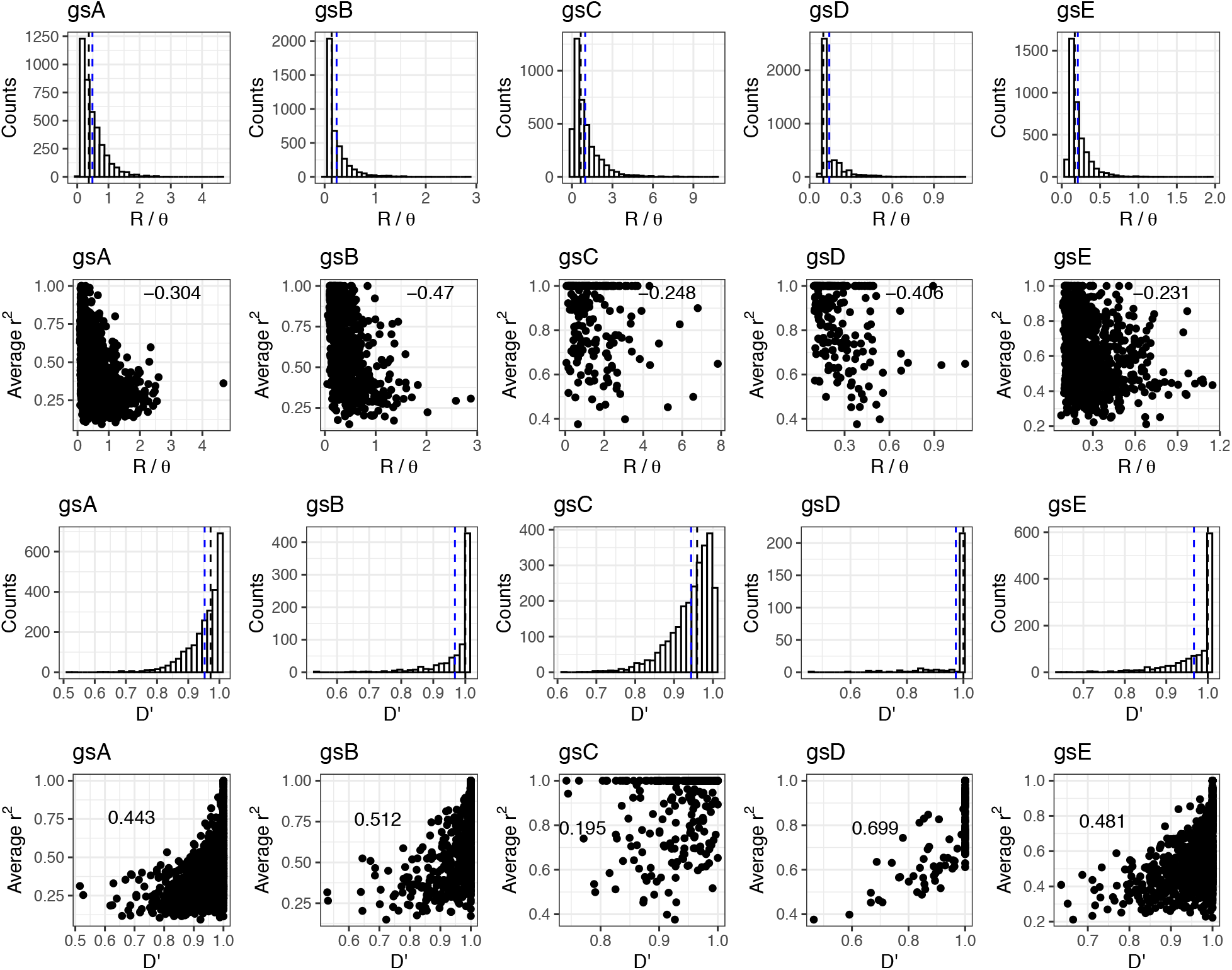
Distribution of R/*θ* and *D* across species and their correlation to *r*^2^. R/*θ* (relative importance of Recombination (R) to mutation (*θ*)) was computed using the software ClonalFrameML (Didelot and Wilson., 2015) by analizing all core orthologs (3086 genes). The distribution of R/*θ* across species is shown (first row), blue and black dashed lines correspond to median and mean R/*θ* values, respectively. The Pearson correlation between R/*θ* and *r*^2^ is shown for each species (second row)). *D′* distributions across species are shown in the third row, and the Pearson’s correlation between *D′* and *r*^2^ metrics are shown in the fourth row.

**Figure S9:**
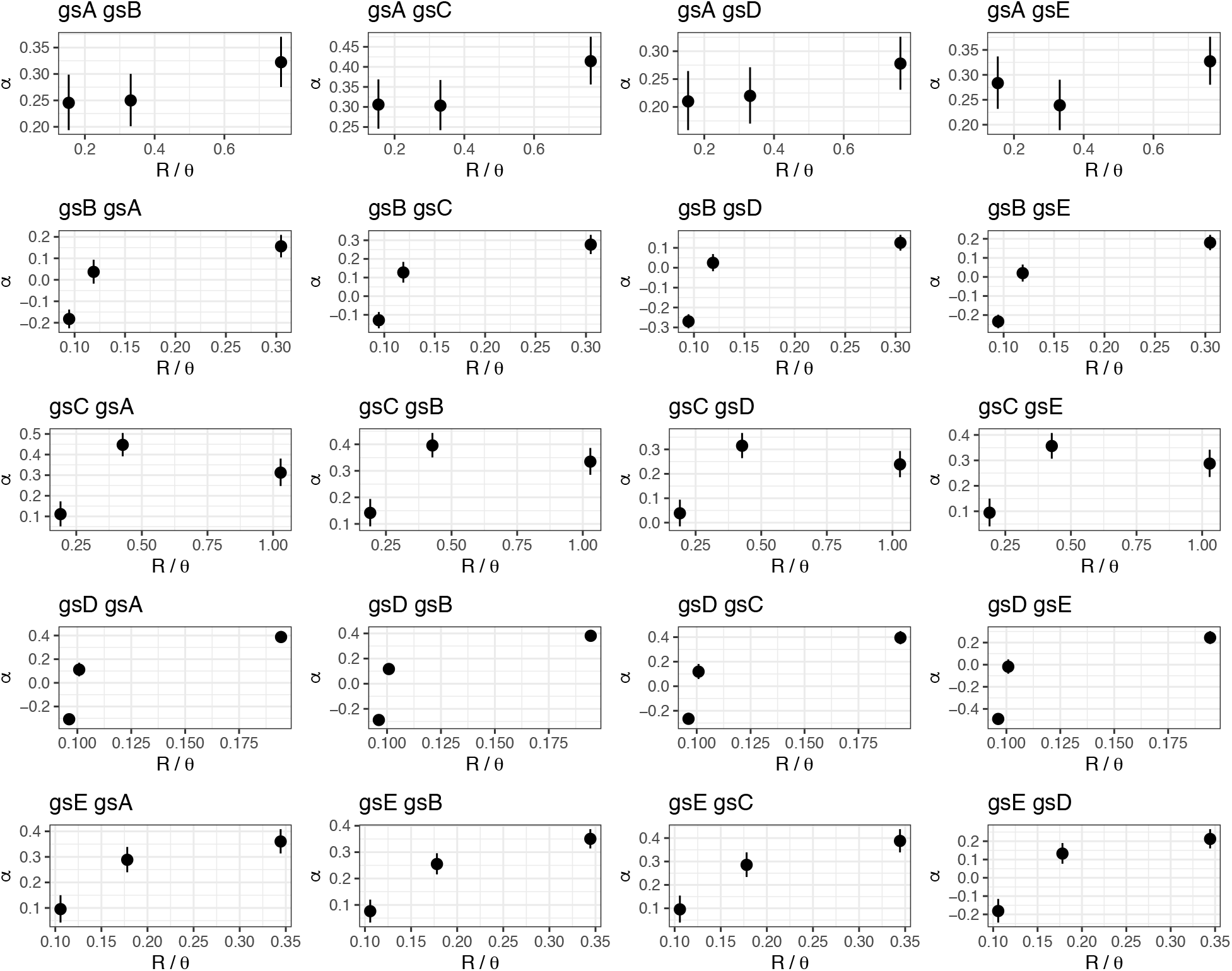
The proportion of adaptive evolution by classes of R/*θ*. The proportion of adaptive evolution (*α*) is computed for each class of recombination (classes of recombination are based on averages values of R/*θ*). For each pairwise estimates of *α* (species1 species2), the polymorphism data from a focal species (species 1) is compared against the divergence counts of an outgroup (species 2), and vice-versa. The *α* estimates (and their associated confidence intervals) were obtained using the best fitting DFE model (GammaZero).

**Figure S10:**
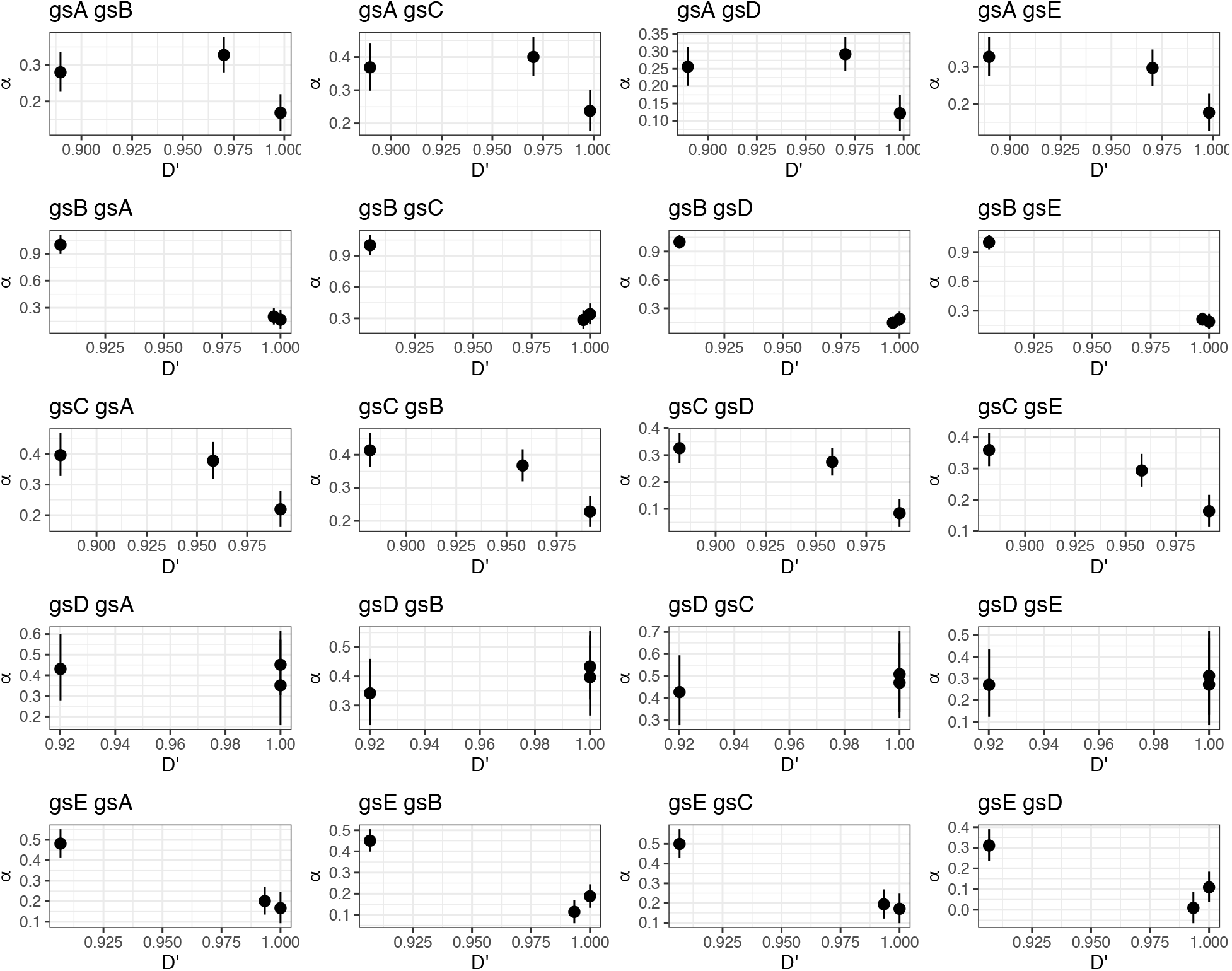
The proportion of adaptive evolution by classes of *D′*. The proportion of adaptive evolution (*α*) is computed for each class of recombination (based on *D′*). For each pairwise estimates of *α* (species1 species2), the polymorphism data from a focal species (species 1) is compared against the divergence counts of an outgroup (species 2), and vice-versa. The *α* estimates (and their associated confidence intervals) were obtained using the best fitting DFE model (GammaZero).

**Figure S11:**
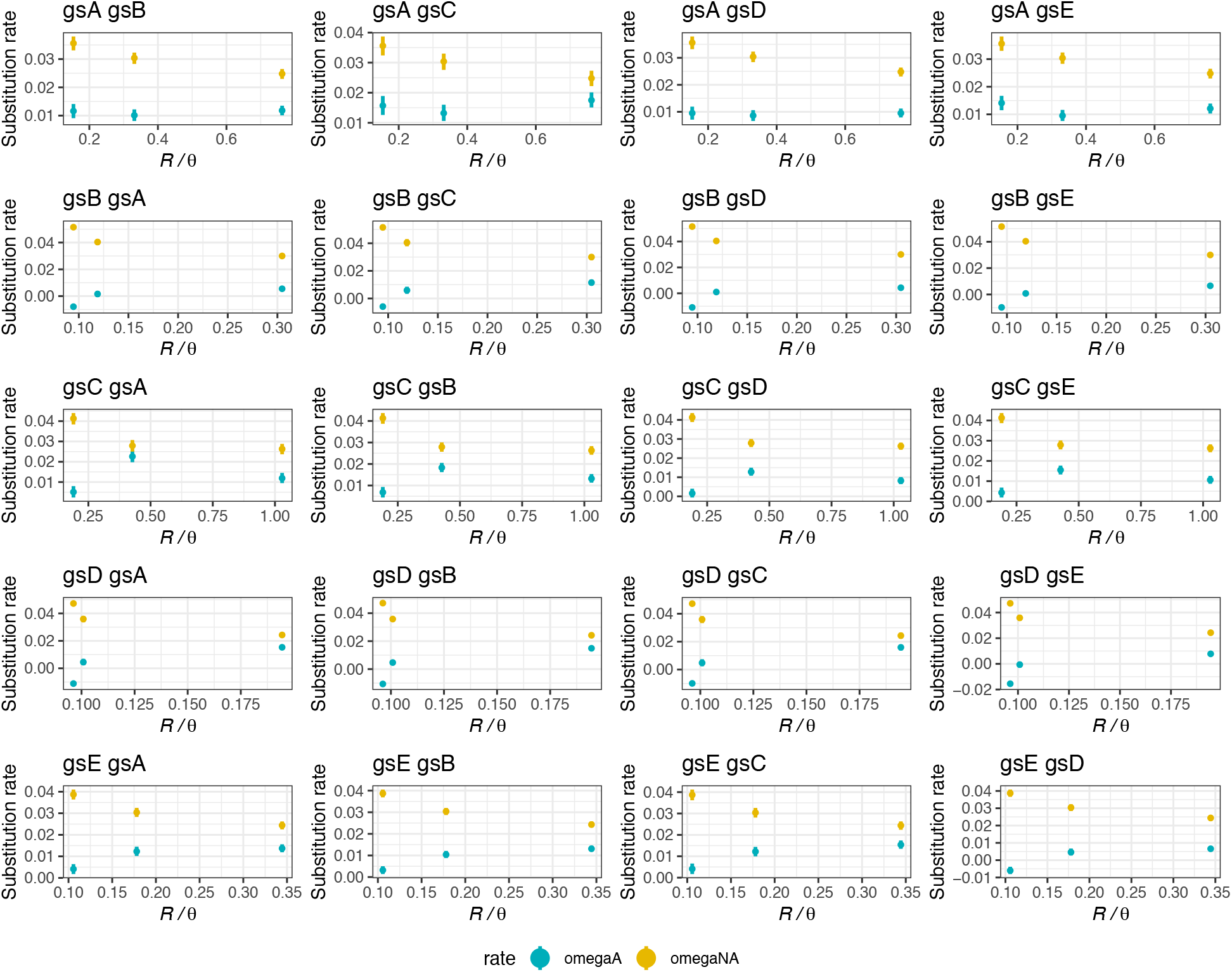
The rate of non-adaptive substitution relative to neutral divergence (*⍵_NA_*), and the rate of adaptive evolution *⍵_A_* were computed for each recombination class (measured by R/*θ*). For each pairwise estimates of *⍵_NA_* and *⍵_A_*, the polymorphism data from one species (species 1) is compared against the divergence counts of an outgroup (species 2), and vice-versa. The *⍵_NA_* and *⍵_A_* estimates (and their associated confidence intervals) were obtained using the GammaZero model.)

**Figure S12:**
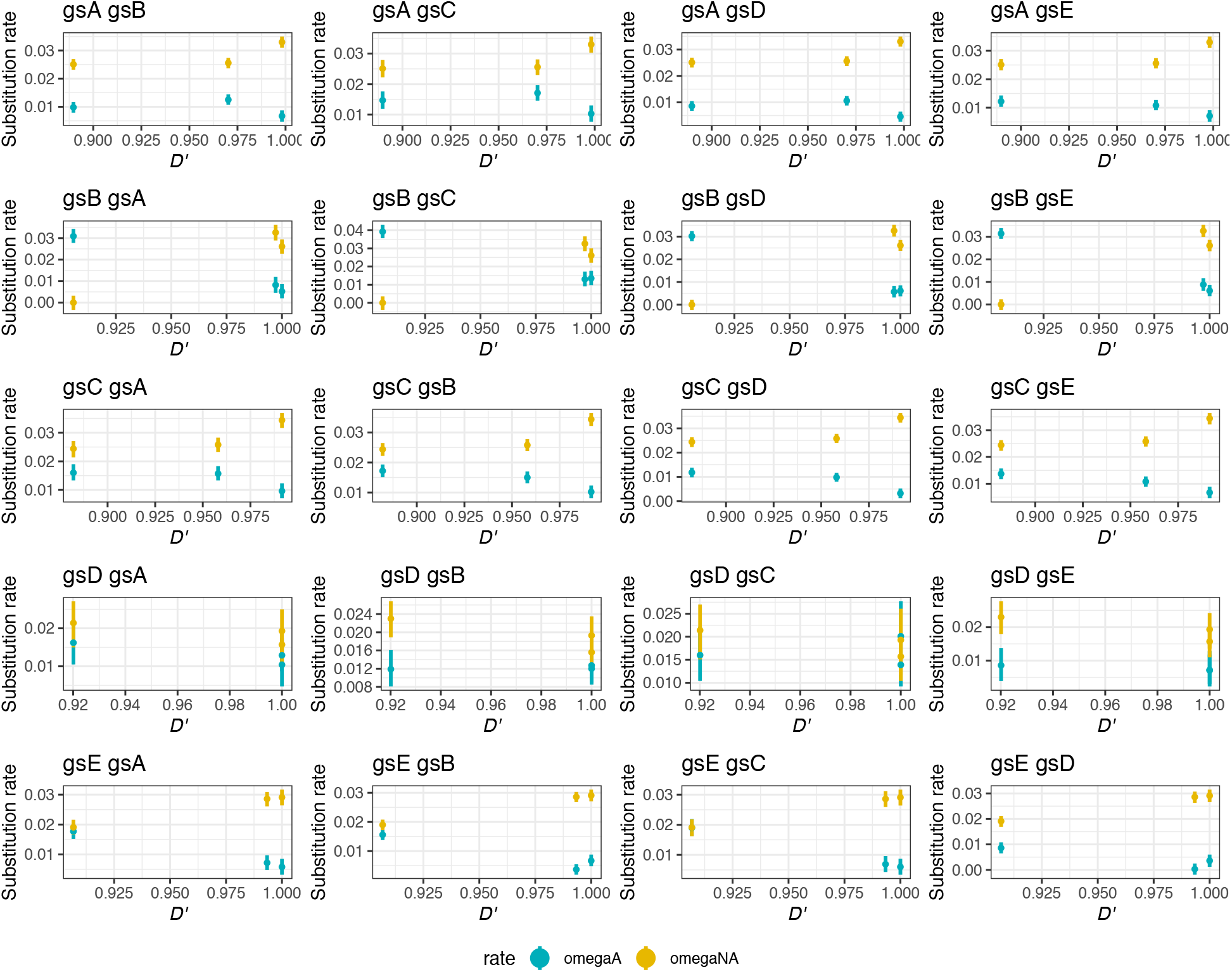
The rate of non-adaptive substitution relative to neutral divergence (*⍵_NA_*), and the rate of adaptive evolution *⍵_A_* were computed for each recombination class (measured by *D′*). For each pairwise estimates of *⍵_NA_* and *⍵_A_*, the polymorphism data from one species (species 1) is compared against the divergence counts of an outgroup (species 2), and vice-versa. The *⍵_NA_* and *⍵_A_* estimates (and their associated confidence intervals) were obtained using the GammaZero model.

**Figure S13:**
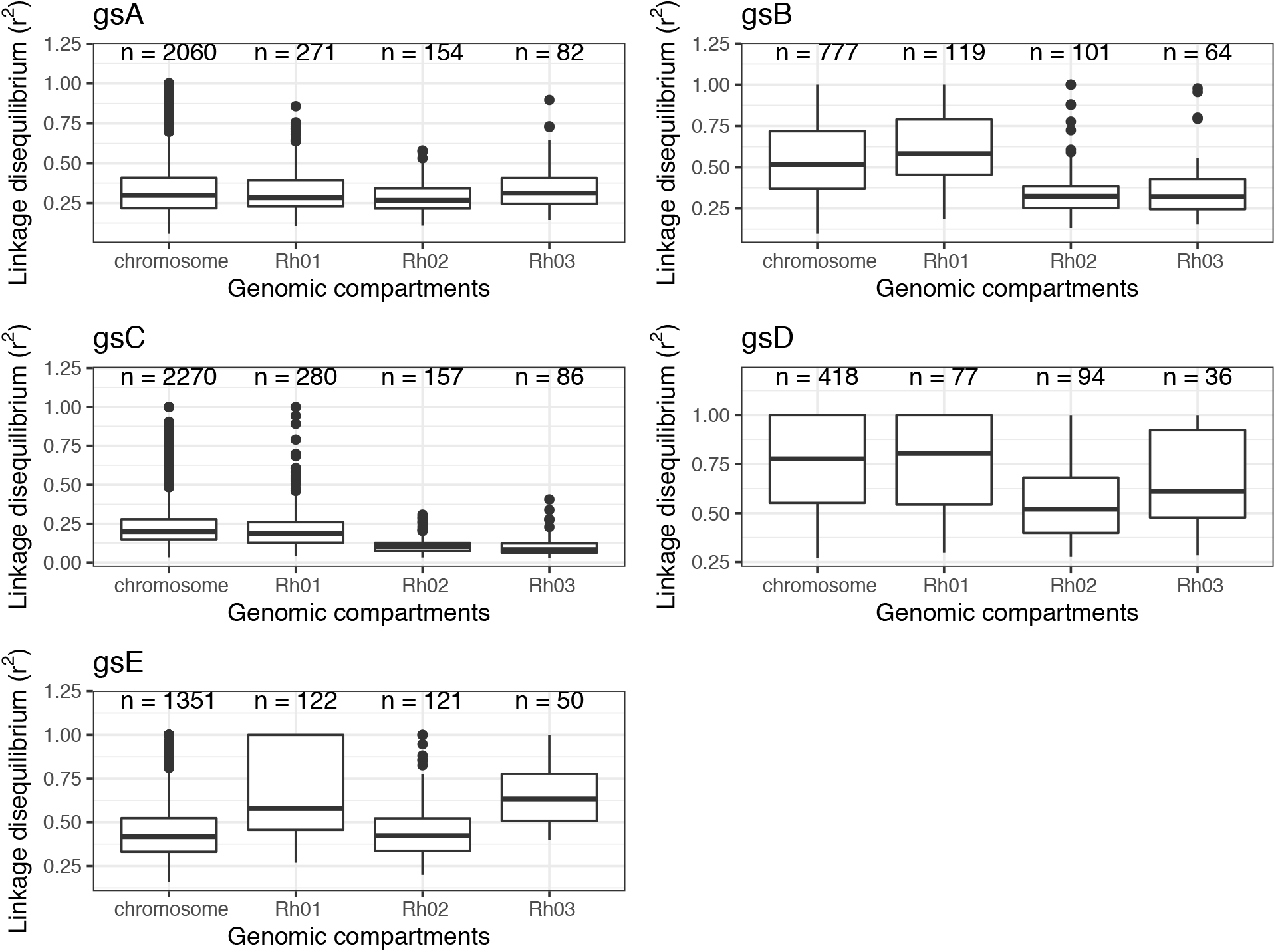
Recombination rate by genomic compartiment. Recombination rate by genomic compartment was comupted based on patterns of linkage disequilibrium (see Methods). Genomic compartiments were previously defined based on sequence variation of a plasmid partitioning gene (repA) that is essential for stable maintenance of nearly all plasmids in Rhizobium (Cavassim et al., 2020). These compartments correspond to the chromosome, chromids (Rh01 and Rh02), and plasmid (Rh03) (Harrison et al., 2010).

**Table S4:** Likelihood differences among DFE models and parameters estimated. Pn and Ps refer to the total number of non-synonymous and synonymous polymorphisms, respectively. Dn and Ds refers to the number of non-synonymous and synonymous substitutions. The total number of non-synonymous and synonymous sites available in the within-species refers to Lpn and Lps, respectively. The number of non-synonymous and synonymous sites observed between-species alignments is denoted by dn and ds, respectively. The models and their corresponding – estimates are shown for a subset of all comparisons.

| pn | Lpn | ps | Lps | dn | ds | Lds | model | X.param | alpha | lnL | AIC | geno |
| --- | --- | --- | --- | --- | --- | --- | --- | --- | --- | --- | --- | --- |
| 18700 | 2413139 | 107889 | 668782.2 | 5993 | 37205 | 668782.2 | Neutral | 61 | -0.076 | -1263.394 | 2648.789 | gsC gsB |
| 18700 | 2413139 | 107889 | 668782.2 | 5120 | 36783 | 668782.2 | GammaZero | 62 | 0.255 | -590.638 | 1305.276 | gsC gsD |
| 18700 | 2413139 | 107889 | 668782.2 | 5120 | 36783 | 668782.2 | ScaledBeta | 63 | 0.736 | -590.177 | 1306.354 | gsC gsD |
| 18700 | 2413139 | 107889 | 668782.2 | 5120 | 36783 | 668782.2 | DisplGamma | 63 | 0.255 | -590.638 | 1307.276 | gsC gsD |
| 18700 | 2413139 | 107889 | 668782.2 | 5120 | 36783 | 668782.2 | GammaGamma | 65 | 0.254 | -590.638 | 1311.277 | gsC gsD |
| 18700 | 2413139 | 107889 | 668782.2 | 5120 | 36783 | 668782.2 | GammaExpo | 64 | 0.243 | -595.904 | 1319.808 | gsC gsD |
| 18700 | 2413139 | 107889 | 668782.2 | 5120 | 36783 | 668782.2 | FGMBesselK | 64 | 0.631 | -714.724 | 1557.448 | gsC gsD |
| 18700 | 2413139 | 107889 | 668782.2 | 5120 | 36783 | 668782.2 | Neutral | 61 | -0.245 | -1263.310 | 2648.620 | gsC gsD |
| 18700 | 2413139 | 107889 | 668782.2 | 5322 | 36086 | 668782.2 | GammaZero | 62 | 0.297 | -590.648 | 1305.296 | gsC gsE |
| 18700 | 2413139 | 107889 | 668782.2 | 5322 | 36086 | 668782.2 | ScaledBeta | 63 | 0.751 | -590.187 | 1306.374 | gsC gsE |
| 18700 | 2413139 | 107889 | 668782.2 | 5322 | 36086 | 668782.2 | DisplGamma | 63 | 0.297 | -590.648 | 1307.296 | gsC gsE |
| 18700 | 2413139 | 107889 | 668782.2 | 5322 | 36086 | 668782.2 | GammaGamma | 65 | 0.296 | -590.648 | 1311.296 | gsC gsE |
| 18700 | 2413139 | 107889 | 668782.2 | 5322 | 36086 | 668782.2 | GammaExpo | 64 | 0.285 | -595.914 | 1319.827 | gsC gsE |
| 18700 | 2413139 | 107889 | 668782.2 | 5322 | 36086 | 668782.2 | FGMBesselK | 64 | 0.652 | -714.734 | 1557.468 | gsC gsE |
| 18700 | 2413139 | 107889 | 668782.2 | 5322 | 36086 | 668782.2 | Neutral | 61 | -0.175 | -1263.320 | 2648.639 | gsC gsE |
| 2325 | 2413139 | 15724 | 668782.2 | 5600 | 40199 | 668782.2 | GammaZero | 6 | 0.251 | -31.135 | 74.270 | gsD gsA |
| 2325 | 2413139 | 15724 | 668782.2 | 5600 | 40199 | 668782.2 | DisplGamma | 7 | 0.256 | -31.127 | 76.255 | gsD gsA |
| 2325 | 2413139 | 15724 | 668782.2 | 5600 | 40199 | 668782.2 | ScaledBeta | 7 | 0.560 | -31.127 | 76.255 | gsD gsA |
| 2325 | 2413139 | 15724 | 668782.2 | 5600 | 40199 | 668782.2 | GammaExpo | 8 | 0.074 | -31.127 | 78.255 | gsD gsA |
| 2325 | 2413139 | 15724 | 668782.2 | 5600 | 40199 | 668782.2 | FGMBesselK | 8 | 0.697 | -31.127 | 78.255 | gsD gsA |
| 2325 | 2413139 | 15724 | 668782.2 | 5600 | 40199 | 668782.2 | GammaGamma | 9 | 0.118 | -31.127 | 80.255 | gsD gsA |
| 2325 | 2413139 | 15724 | 668782.2 | 5600 | 40199 | 668782.2 | Neutral | 5 | -0.061 | -41.088 | 92.175 | gsD gsA |
| 2325 | 2413139 | 15724 | 668782.2 | 9790 | 70104 | 668782.2 | GammaZero | 6 | 0.254 | -31.693 | 75.385 | gsD gsB |
| 2325 | 2413139 | 15724 | 668782.2 | 9790 | 70104 | 668782.2 | DisplGamma | 7 | 0.258 | -31.685 | 77.369 | gsD gsB |
| 2325 | 2413139 | 15724 | 668782.2 | 9790 | 70104 | 668782.2 | ScaledBeta | 7 | 0.564 | -31.685 | 77.369 | gsD gsB |
| 2325 | 2413139 | 15724 | 668782.2 | 9790 | 70104 | 668782.2 | GammaExpo | 8 | 0.062 | -31.685 | 79.369 | gsD gsB |
| 2325 | 2413139 | 15724 | 668782.2 | 9790 | 70104 | 668782.2 | FGMBesselK | 8 | 0.697 | -31.685 | 79.369 | gsD gsB |
| 2325 | 2413139 | 15724 | 668782.2 | 9790 | 70104 | 668782.2 | GammaGamma | 9 | 0.107 | -31.685 | 81.369 | gsD gsB |
| 2325 | 2413139 | 15724 | 668782.2 | 9790 | 70104 | 668782.2 | Neutral | 5 | -0.059 | -41.645 | 93.290 | gsD gsB |
| 2325 | 2413139 | 15724 | 668782.2 | 5120 | 36783 | 668782.2 | GammaZero | 6 | 0.250 | -31.046 | 74.092 | gsD gsC |
| 2325 | 2413139 | 15724 | 668782.2 | 5120 | 36783 | 668782.2 | DisplGamma | 7 | 0.255 | -31.038 | 76.076 | gsD gsC |
| 2325 | 2413139 | 15724 | 668782.2 | 5120 | 36783 | 668782.2 | ScaledBeta | 7 | 0.560 | -31.038 | 76.076 | gsD gsC |
| 2325 | 2413139 | 15724 | 668782.2 | 5120 | 36783 | 668782.2 | GammaExpo | 8 | 0.076 | -31.038 | 78.076 | gsD gsC |
| 2325 | 2413139 | 15724 | 668782.2 | 5120 | 36783 | 668782.2 | FGMBesselK | 8 | 0.697 | -31.038 | 78.076 | gsD gsC |
| 2325 | 2413139 | 15724 | 668782.2 | 5120 | 36783 | 668782.2 | GammaGamma | 9 | 0.120 | -31.038 | 80.076 | gsD gsC |
| 2325 | 2413139 | 15724 | 668782.2 | 5120 | 36783 | 668782.2 | Neutral | 5 | -0.062 | -40.998 | 91.997 | gsD gsC |
| 2325 | 2413139 | 15724 | 668782.2 | 4713 | 39531 | 668782.2 | GammaZero | 6 | 0.125 | -31.041 | 74.081 | gsD gsE |
| 2325 | 2413139 | 15724 | 668782.2 | 4713 | 39531 | 668782.2 | DisplGamma | 7 | 0.130 | -31.033 | 76.065 | gsD gsE |
| 2325 | 2413139 | 15724 | 668782.2 | 4713 | 39531 | 668782.2 | ScaledBeta | 7 | 0.486 | -31.033 | 76.065 | gsD gsE |
| 2325 | 2413139 | 15724 | 668782.2 | 4713 | 39531 | 668782.2 | GammaExpo | 8 | -0.082 | -31.033 | 78.065 | gsD gsE |
| 2325 | 2413139 | 15724 | 668782.2 | 4713 | 39531 | 668782.2 | FGMBesselK | 8 | 0.646 | -31.033 | 78.065 | gsD gsE |
| 2325 | 2413139 | 15724 | 668782.2 | 4713 | 39531 | 668782.2 | GammaGamma | 9 | -0.030 | -31.033 | 80.065 | gsD gsE |
| 2325 | 2413139 | 15724 | 668782.2 | 4713 | 39531 | 668782.2 | Neutral | 5 | -0.240 | -40.993 | 91.986 | gsD gsE |
| 5069 | 2413139 | 37227 | 668782.2 | 5764 | 39059 | 668782.2 | GammaZero | 9 | 0.270 | -63.961 | 145.922 | gsE gsA |
| 5069 | 2413139 | 37227 | 668782.2 | 5764 | 39059 | 668782.2 | DisplGamma | 10 | 0.270 | -63.961 | 147.922 | gsE gsA |
| 5069 | 2413139 | 37227 | 668782.2 | 5764 | 39059 | 668782.2 | ScaledBeta | 10 | 0.374 | -64.157 | 148.315 | gsE gsA |
| 5069 | 2413139 | 37227 | 668782.2 | 5764 | 39059 | 668782.2 | GammaExpo | 11 | 0.232 | -63.892 | 149.784 | gsE gsA |
| 5069 | 2413139 | 37227 | 668782.2 | 5764 | 39059 | 668782.2 | FGMBesselK | 11 | 0.600 | -64.455 | 150.910 | gsE gsA |
| 5069 | 2413139 | 37227 | 668782.2 | 5764 | 39059 | 668782.2 | GammaGamma | 12 | 0.868 | -63.896 | 151.792 | gsE gsA |
| 5069 | 2413139 | 37227 | 668782.2 | 5764 | 39059 | 668782.2 | Neutral | 8 | 0.077 | -86.821 | 189.642 | gsE gsA |
| 5069 | 2413139 | 37227 | 668782.2 | 9536 | 66442 | 668782.2 | GammaZero | 9 | 0.250 | -64.478 | 146.957 | gsE gsB |
| 5069 | 2413139 | 37227 | 668782.2 | 9536 | 66442 | 668782.2 | DisplGamma | 10 | 0.250 | -64.478 | 148.957 | gsE gsB |
| 5069 | 2413139 | 37227 | 668782.2 | 9536 | 66442 | 668782.2 | ScaledBeta | 10 | 0.356 | -64.675 | 149.349 | gsE gsB |
| 5069 | 2413139 | 37227 | 668782.2 | 9536 | 66442 | 668782.2 | GammaExpo | 11 | 0.205 | -64.409 | 150.819 | gsE gsB |
| 5069 | 2413139 | 37227 | 668782.2 | 9536 | 66442 | 668782.2 | FGMBesselK | 11 | 0.588 | -64.973 | 151.945 | gsE gsB |
| 5069 | 2413139 | 37227 | 668782.2 | 9536 | 66442 | 668782.2 | GammaGamma | 12 | 0.864 | -64.413 | 152.826 | gsE gsB |
| 5069 | 2413139 | 37227 | 668782.2 | 9536 | 66442 | 668782.2 | Neutral | 8 | 0.051 | -87.338 | 190.677 | gsE gsB |
| 5069 | 2413139 | 37227 | 668782.2 | 5322 | 36086 | 668782.2 | GammaZero | 9 | 0.270 | -63.882 | 145.763 | gsE gsC |
| 5069 | 2413139 | 37227 | 668782.2 | 5322 | 36086 | 668782.2 | DisplGamma | 10 | 0.270 | -63.882 | 147.763 | gsE gsC |
| 5069 | 2413139 | 37227 | 668782.2 | 5322 | 36086 | 668782.2 | ScaledBeta | 10 | 0.374 | -64.078 | 148.156 | gsE gsC |
| 5069 | 2413139 | 37227 | 668782.2 | 5322 | 36086 | 668782.2 | GammaExpo | 11 | 0.233 | -63.813 | 149.625 | gsE gsC |

